# A prefrontal cortex-hypothalamus circuit for heart rate control in non-human primates

**DOI:** 10.64898/2026.09.18.752614

**Authors:** Kevin Xu, Takaya Ogasawara, Jinyun Yuan, Ethan S. Bromberg-Martin, Mengxi Yun, Steven P. Errington, Andrew H. Stark, Takafumi Minamimoto, Ken-ichi Inoue, Hong Chen, Ilya E. Monosov

## Abstract

Humans exhibit changes in heart rate during cognitive and emotional events, and dysregulation of this brain–heart coupling is a hallmark of many psychiatric disorders. Yet, how circuits that regulate cognition exert direct control over heart physiology remains poorly understood, particularly in primates. Here, we identify a prefrontal–hypothalamic circuit that exerts powerful, bidirectional control over heart rate in monkeys. We show that electrical stimulation of a small ventral subregion within the ventrolateral prefrontal cortex (area 47/12; a12) evokes rapid and robust decreases in heart rate and increases heart rate variability, whereas neighboring prefrontal regions produce weaker or no effects. And a12 neurons are coupled to heart rate on a moment-by-moment basis. Anatomical tracing data reveal projections from a12 to a circumscribed region of lateral hypothalamus (LHA), and stimulation of this LHA target recapitulates the cardiac effects of prefrontal activation. Consistent with a prefrontal-to-hypothalamus system for heart rate control, stimulation of LHA produces faster, stronger, state-independent heart effects, whereas stimulation of a12 produces effects that are sensitive to internal states. Oppositely to its activation, chemogenetic inactivation of a12 or its projections to LHA increases heart rate, demonstrating that this pathway can provide tonic inhibitory control over cardiac function. To make steps towards a more cell-type-specific and minimally invasive causal control of this circuit, we developed a primate sonogenetic approach based on ultrasound activation of CaMKII+ neurons virally transduced with TRPV1. Sonogenetic stimulation of a12 produced increases in heart rate variability, and these effects were attenuated by inactivation of LHA. Finally, we found that sonogenetic and chemogenetic perturbation of the same prefrontal region during a value-based decision-making task bidirectionally alters risky decision making. Together, our work identifies how a12 regulates heart rate and cognition, revealing a mechanistic substrate for brain–body interactions and a potential therapeutic target for disorders of cognitive–autonomic dysregulation.

## INTRODUCTION

Humans experience rapid and profound changes in heart rate during cognitive and emotional events. Common expressions such as “you broke my heart” or “my heart just skipped a beat” reflect common intuitions about the powerful brain-heart relationship. Indeed, dysregulation of this brain–heart coupling is a hallmark of many psychiatric disorders (*1*, *2*). While we know from work across many animal models that a distributed set of mammalian brain areas referred to as the central autonomic network (CAN) can impact heart rate (*3–8*), the neural pathways and mechanisms by which our cognition exerts direct, causal control over moment-to-moment cardiac dynamics remain poorly understood, especially in the primate brain.

Understanding how heart rate dynamics are controlled by prefrontal circuits is critical not only for basic neuroscience but also for the clinic. Heart rate dynamics provide accessible, non-invasive readouts of internal physiological state (*9*, *10*) and are strongly associated with emotional regulation (*11*), cognitive control (*12–16*), and mental health outcomes (*17–19*). In many psychiatric disorders, cardiac dynamics are poorly aligned with task demands and ongoing cognitive operations, suggesting that brain systems governing cognition may fail to appropriately regulate or coordinate bodily states, including heart rate. However, these issues remain largely unstudied at the level of identified circuits in the primate brain, the model whose prefrontal organization most closely resembles that of humans.

Older studies involving large lesions (*20*, *21*), electrical stimulation trains known to produce gross effects (*22*, *23*), and relatively recent neuroimaging (*24–26*) together suggest that the primate prefrontal cortex could be an important locus of top-down control over cardiac dynamics. However, how and through which circuits this control is exerted remains to be causally and mechanistically established.

Here, using modern and novel circuit perturbation approaches and computational analyses of heart rate dynamics, neural activity, and behavior, we identify a prefrontal–hypothalamic circuit in macaques that provides direct, causal, and hierarchical control over heart rate, and links heart related prefrontal circuitry to decision-making. We show that a circumscribed region of the ventrolateral prefrontal cortex, area 47/12 (a12), strongly influences cardiac state via its projections to lateral hypothalamus, and that this pathway exerts bidirectional control over heart rate—activation slows the heart, whereas inactivating the pathway accelerates it. Single a12 neurons’ activity track, and can precede, spontaneous changes in heart rate. Within the a12-hypothalamus circuit, the hypothalamus serves as a fast effector-controller that is largely state-independent, whereas the prefrontal cortex provides slower, relatively state-dependent top-down control in part through the hypothalamus. Leveraging our findings, we developed a relatively cell-type-specific, minimally invasive sonogenetic approach to perturb a12 and modulate heart rate dynamics. Finally, we find that perturbing a12 bidirectionally alters risky choice behavior and show how a12 links cognition and cardiac state while keeping their neural codes largely separable, positioning it to coordinate the two flexibly in a context dependent manner.

## RESULTS

### Identification of a prefrontal–hypothalamic pathway for rapid control of heart rate

We sought to identify a pathway through which the prefrontal cortex (PFC) could exert rapid control over cardiac dynamics in non-human primates beginning with a bottom-up, circuit-discovery approach. We start by focusing on the hypothalamus, a major recipient of topographic prefrontal projections and a key autonomic control hub with dense outputs to brainstem autonomic nuclei that control cardiac function (*21*, *27*, *28*). We first mapped cardiac responses to focal electrical stimulation across multiple subregions of the primate hypothalamus, and later verified these results with chemogenetics and/or pharmacology. Focal electrical stimulation (Methods) remains the only technique that would allow us to functionally map large swaths of the primate hypothalamus, without strong anatomical bias. This initial mapping revealed a localized region within the lateral hypothalamus (LHA) whose stimulation rapidly suppressed heart rate. We then used this region as an entry point to examine whether and how the prefrontal cortex can regulate heart rate dynamics through the hypothalamus (**Fig. 1a**) and verified all our findings with additional modern circuit perturbation methods.

**Figure 1:**
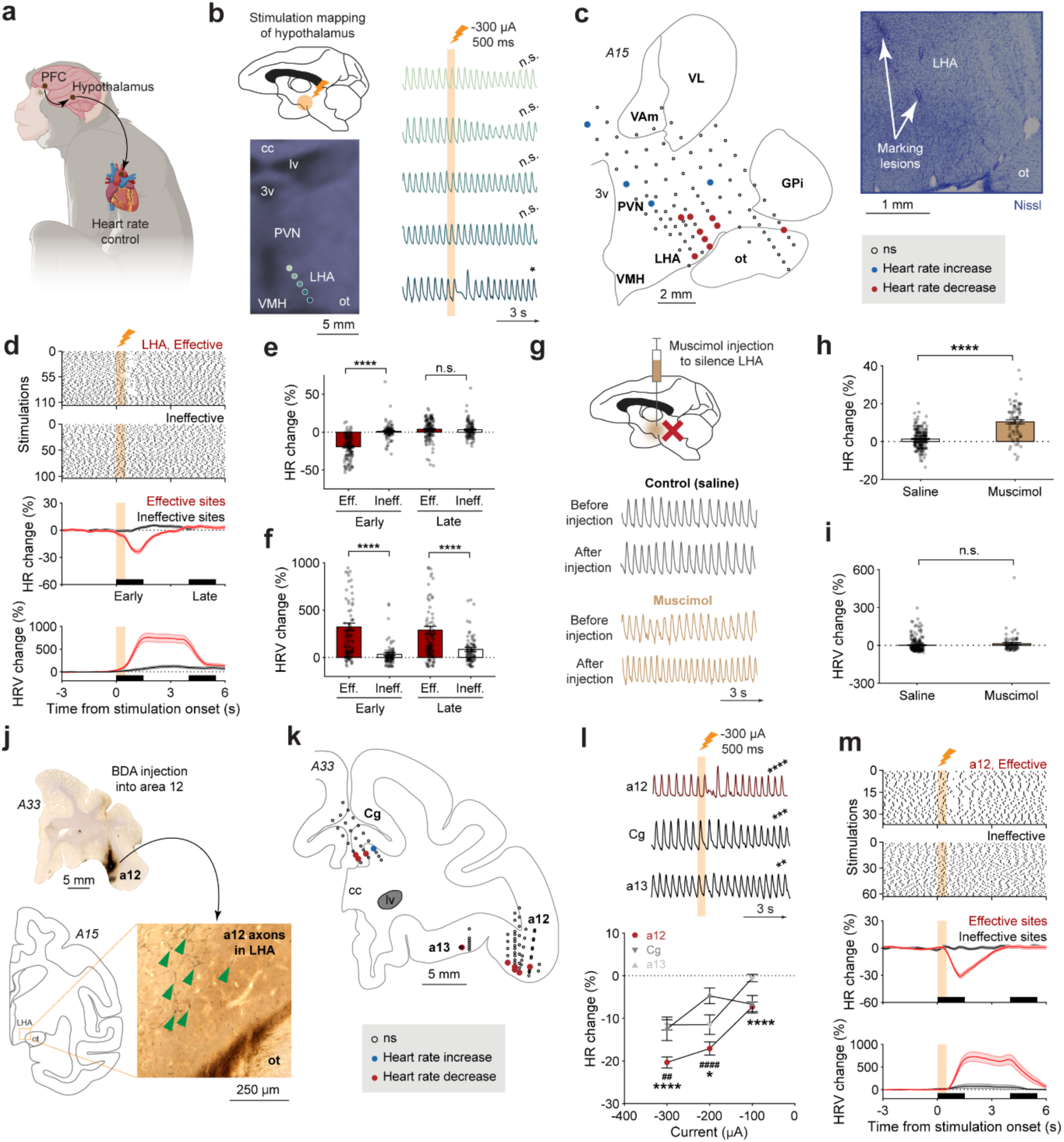
Identification of a prefrontal–hypothalamic pathway for rapid influence over cardiac dynamics. a, Schematic illustration of the putative primate PFC-hypothalamus system that we study in the context of heart rate modulation. **b,** Stimulation mapping of hypothalamus. Left: T1-weighted MRI showing stimulation sites near LHA (cc, corpus callosum; lv, lateral ventricle; 3v, third ventricle; PVN, paraventricular nucleus; VMH, ventromedial hypothalamus; LHA, lateral hypothalamus). Right: Representative photoplethysmograph (PPG) traces as a function on depth in LHA. Yellow bar indicates electrical stimulation for 500 ms (−300 μA). Asterisks indicate significant decrease in the heart rate averaged 0-1.5 s after stimulation onset. n = 7, 5, 5, 5,10 stimulations as a function of depth (dorsal to ventral); P = 0.3125, 0.4375, 0.3125, 0.0625, 0.0371 (two-tailed signed-rank test against a hypothetical median of 0). **c,** Reconstruction of recording sites in LHA. Left: Schematic illustration of monkey brain in the coronal plane at interaural A15. Circles indicate stimulation sites. Red indicates significant heart rate decrease, blue indicates significant heart rate increase, and un-filled indicates no significant heart rate change (significance defined as p < 0.05, signed-rank test against a hypothetical median of 0). Right: Histological image of electrolytic lesion marker in LHA confirming the stimulation site. (VAm, magnocelluar division of ventral anterior thalamic nucleus; VL, ventral lateral thalamic nucleus; GPi, globus pallidus internal; ot, optic tract). **d,** Heart rate time course in LHA during electrical stimulation. Yellow bar indicates duration of electrical stimulation (500 ms; −300 μA). Top: Raster of the heartbeats in effective and ineffective sites within LHA during electrical stimulation. Middle: Average heart rate change time course for effective (red) and ineffective (grey) sites. Shaded regions represent s.e.m. Black bars on the x-axis correspond to the early (0-1.5 s) and late (4-5.5 s) time windows. Bottom: Average heart rate variability changes time course for effective (red) and ineffective (grey) sites. **e,** Average heart rate changes evoked by effective and ineffective site stimulation, averaged in early and late windows. Data are mean ± s.e.m. (error bars too small to see); n = 116, 100 stimulations for effective and ineffective sites, respectively. P = 2.26e-21, 0.572 for early and late windows, respectively (two-tailed rank-sum test). **f,** Heart rate variability changes evoked by effective and ineffective site stimulation, averaged in early and late windows. Data are mean ± s.e.m. (error bars too small to see). P = 1.09e-13, 8.21e-06 for early and late windows, respectively (two-tailed rank-sum test). **g,** Muscimol injection to silence LHA. Top, PPG traces before and after injection of saline. Bottom, PPG traces before and after injection of muscimol. **h,** Average heart rate change after injection of saline or muscimol. Data are mean ± s.e.m. (too small to see); n = 217, 75 measurements for saline and muscimol, respectively. P = 1.72e-23 (two-tailed t-test). **i,** Average heart rate variability changes after injection of saline or muscimol. Data are mean ± s.e.m. (too small to see). P = 0.158 (two-tailed t-test). **j,** Top: Histological image of BDA injection into area 12. Bottom: Histological image of BDA projection sites within LHA. Green arrows highlight the a12 axons observed in LHA. **k,** Reconstruction of recording sites in the prefrontal cortex. Left, schematic illustration of monkey brain in the coronal plane at interaural A33. Circles follow the same convention as (c). Right, representative PPG traces in areas 12, Cg, and 13. (Cg, cingulate). **l,** Top: Representative PPG traces for each prefrontal cortex region. Asterisks indicate significant decrease in the heart rate for each site in the early window. P = 7.11e-13, 2.61e-08, 9.94e-05 corresponding to a12, Cg, a13, respectively (signed-rank test against a hypothetical median of 0). Bottom: Average heart rate changes in areas 12 (red), Cg (dark grey), and 13 (light grey) as a function of current. Data are mean ± s.e.m. * denotes significance comparison between a12 and a13, # denotes comparison between a12 and Cg. For a12, n = 72, 46, 22 stimulations for −300, −200, −100 μA, respectively; for Cg, n = 41, 15, 20 stimulations; for a13, n = 27, 32, 10 stimulations. For a12 vs. Cg, P = 2.72e-06, 0.0436, 3.07e-05 for −300, −200, −100 μA, respectively; for a12 vs a13, P = 0.00122, 4.34e-05, 0.951 (two-tailed rank-sum test comparing a12 vs. Cg or a12 vs. a13.). **m,** Heart rate time course in a12 during electrical stimulation, following the same format as (d). Throughout this figure, *, **, ***, **** indicate P < 0.05, 0.01, 0.001, 0.0001, respectively; n.s., non-significant.

We found that stimulation-evoked suppressive effects on heart rate were highly localized within a subregion of the hypothalamus. Specifically, stimulation of a particular area within LHA, often in the region above the optic tract, rapidly evoked a marked decrease in heart rate, akin to the heart skipping a beat (**Fig. 1b–c**; **Fig. S2** for additional reconstruction verification; **Fig. S3a-c**). In contrast, stimulation of more dorsal sites along the same electrode trajectories did not evoke significant short timescale changes in heart rate. The effects were not artifactual or signal dropouts and were associated with relatively large and reliable effects (**Fig. 1-2**; **Fig. S1b, c**). Also, these heart rate effects were not simply a consequence that followed overt behavioral responses evoked by stimulation (**Fig. S4**). In addition, we found that heart rate decrease effects were readily observable upon on-target stimulation in LHA even when the animal’s eyes were closed and remained closed well after stimulation was over (**Fig. S5**).

**Figure 2:**
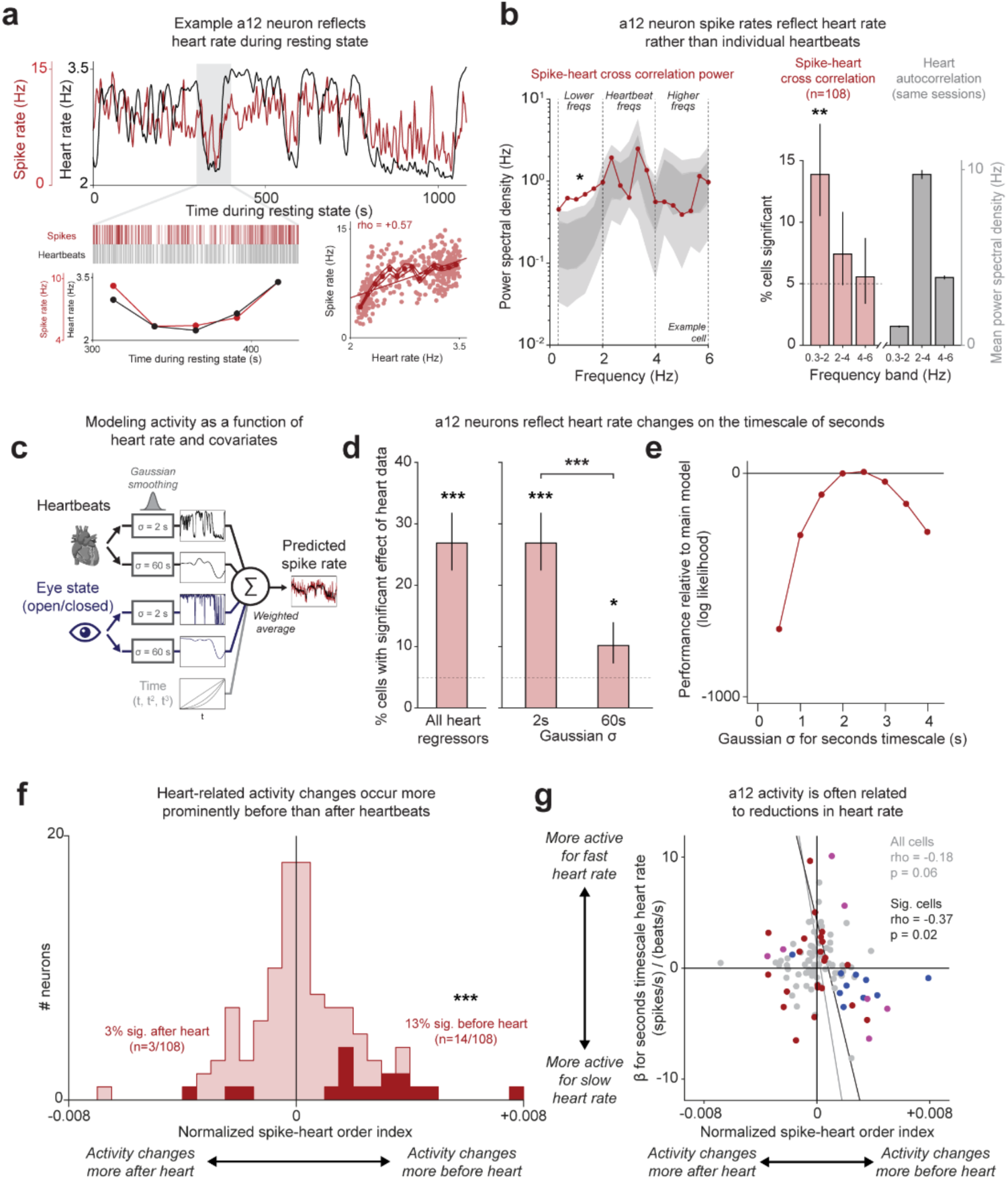
a12 neuronal activity tracks spontaneous fluctuations in heart rate. **a**, An example a12 neuron during resting state session with spiking activity (red, smoothed with Gaussian kernel σ = 2 s) correlated with heart rate (black, same). Bottom left: detailed view of correlated changes in spike and heartbeat trains (red and black; dots are mean rates in 20 s bins, error bars represent s.e.m., too small to see). Bottom right: spike rate vs. heart rate correlation (quantified using 3 s time bins; text indicates rank correlation). **b**, a12 neurons reflect heart rate rather than individual heartbeats. Left: Power spectral density of cross-correlation between spike train and heartbeat train for the example neuron (red) compared to n=100 randomly time-shifted controls representing the null hypothesis that spikes are unrelated to heartbeats (dark and light gray are central 68% and 95% of the control distribution). * indicates the mean power spectral density in a frequency band is significantly different between real vs. controls (p < 0.05, two-tailed circular shift tests). Right: % of a12 neurons with significantly different power spectral density of spike-heart cross-correlation in each frequency band (red), which only occurs for lower frequency band (0.3-2 Hz). ** indicates above chance (p < 0.01, one-tailed binomial test). By contrast, the mean power spectral density for heartbeat autocorrelation in the same frequency bands (gray) is greatest in the heartbeat frequency band (2-4 Hz). **c**, Model predicting spike rate at each millisecond (right, red vs. black are real vs predicted rates) as a weighted combination of heart rate (top, black) smoothed on the order of seconds (σ = 2 s) and minutes (σ = 60 s), eye open/closed state smoothed the same way (middle, blue), and linear, quadratic, and cubic effects of time (bottom, gray). **d**, Left: % of neurons whose fits are significantly improved by including the real heartbeat data, compared to n=100 randomly time-shifted controls. *** indicates above chance (p < 0.001, one-tailed binomial test). Right: % of neurons with significant fitted weights of heart rate smoothed at each timescale. *, *** indicates above chance (p < 0.05, 0.001, one-tailed binomial tests). More neurons have significant weights on the timescale of seconds (p < 0.001, exact test of proportions). **e**, Relative log likelihood of fitted models using different σ for the seconds timescale (red). The best fit is σ = 2-2.5 s. **f**, histogram of each neuron’s normalized spike-heart order index indicating whether a neuron’s spike-heart cross-correlation deviates from chance significantly more before heartbeats (index > 0) or after heartbeats (index < 0) (p < 0.05, circular shift test). More neurons than expected by chance have indexes > 0 (p < 0.001, binomial test). **g**, Relationship between each neuron’s normalized spike-heart order index (x-axis) and fitted weight in the model above for seconds timescale effect of heart rate (y-axis). Blue, red, and purple indicate cells significant along x, y, and both axes. Lines indicate results of type 2 linear regression and text indicates rank correlation and its p-value, for all cells (gray) and all cells for which x or y is significant (black).

We also observed several anatomically distributed sites within the dorsal hypothalamus that had the opposite effects on cardiac rhythms, effectively increasing rather than decreasing heart rate, but these effects were weaker and occurred on distinct slower timescales (**Fig. 1c, Fig. S2c-e**).

To visualize the cross-validated effects of LHA stimulation, we classified each site as producing a significant decrease in heart rate (“effective sites”) or as producing no detectable effect (“ineffective sites”), visualized the responses in one half of the stimulation trials, and performed statistical analysis of the heart responses in the other half of stimulation trials. This analysis revealed a rapid, reliable, and highly stereotyped heart rate decrease for effective sites within LHA that was absent for ineffective sites (**Fig. 1d, top**). The corresponding average time courses show an immediate decrease in heart rate following stimulation onset (**Fig. 1d**). We quantified these heart rate effects using two time windows: an early window (0–1.5 s after stimulation onset) and a late window (4–5.5 s after stimulation onset). On average, effective site stimulation evoked a heart rate change of −21.0 ± 0.1% in the early window and a weaker tendency of +4.3 ± 0.1% in the late window (**Fig. 1d**). The heart rate decrease effect in the early window was significantly different from ineffective site stimulation (**Fig. 1e**). We also examined heart rate variability (HRV), another marker reflecting the autonomic nervous system’s ability to regulate heart function (*9*). In part related to the rate changes, stimulation of the “effective sites” was associated with marked increases in HRV in both early and late time windows (360 ± 4% and 349 ± 4%, respectively) (**Fig. 1f**).

We next performed pharmacological knockdowns of the effective sites within LHA with muscimol injections. Muscimol inactivates local neurons by agonizing their GABA-A receptors, while sparing fibers of passage (*29*, *30*). We found that muscimol injections into LHA produced significant tonic heart rate *increase* effects (**Fig. 1g-i**), opposite to the *decrease* we observed following stimulation.

In sum, electrical stimulation and muscimol injection experiments identified a region within LHA whose focal causal manipulation is sufficient to produce consistent perturbations of cardiac dynamics.

We next asked whether and how the PFC can access the heart-related regions of the LHA. Prior anatomical studies indicate that the LHA receives inputs from several PFC areas involved in decision making and cognition, with most prominent projections arising from area 12 (lateral orbitofrontal cortex), but also from areas 13 (central orbitofrontal cortex), and 24 (cingulate cortex) (*31–35*). Though relatively anatomically and functionally underexplored, area 12 has recently been implicated in cognitive-control and decision-making under uncertainty and risk, states which have important implications for cardiac function (*36–39*). It has also emerged as a target of interest in psychiatric disorders in which autonomic responses are prominently disrupted (*40*).

We (i) tested whether projections from area 12 to LHA overlap the cardiac-effective LHA subregion identified above (**Fig. 1c**), (ii) compared the cardiac effects of stimulation in area 12 with those evoked from PFC areas 13 and 24 that also project to LHA to determine whether area 12 provides a particularly potent prefrontal access point to this cardiac control system, and (iii) tested if PFC and LHA influences on the heart are consistent with a hierarchical control system, such that PFC has a slower yet flexible influence modulated by the current state of the animal while LHA influence is rapid and more context-independent.

We analyzed an injection experiment in which biotinylated dextran amine (BDA) was injected into a12. Consistent with an a12 to LHA projection, we visualized rich axon and axon-bouton staining in LHA (**Fig. 1j**, Methods). Crucially, axons were found in the region of the LHA which was particularly enriched with cardiac-related sites in our stimulation experiments (**Fig. 1c**; *additional cases support similar conclusions and are available upon request*).

We next used the same electrical stimulation paradigm as in **Fig. 1b** to map heart rate responses within a12, a13, and the cingulate cortex. Like LHA, electrical stimulation of some sites within all 3 regions evoked a decrease in heart rate (**Fig. 1k**). However, by comparing the effective site responses among the three cortical areas (effective sites defined as in **Fig. 1d**), we observed that a12 often evoked the greatest decrease in heart rate (**Fig. 1l**). We interpret these results to mean that while each area may contain sites that can regulate heart rate dynamics, a12 regions that project to the LHA, in particular, have highly pronounced effects on cardiac dynamics.

a12 stimulation resulted in a rapid and reliable heart rate decrease for effective sites with similar decrease and rebound kinetics compared with LHA stimulation (**Fig. 1m, top**). Effective-site a12 stimulation evoked a heart rate change of −21.6 ± 0.2% in the early window and +0.7 ± 0.3% in the late window, the former of which was significantly different from ineffective site stimulation (**Fig. S3d-e**). For heart rate variability, similar to LHA stimulation, stimulation of effective sites in a12 also had a significantly greater effect than stimulation of ineffective sites in the early and late time windows (213 ± 5%, 403 ± 14%, respectively) (**Fig. S3f**).

### Neural activity in area 12 reflects heart activity

Having found that a12 stimulation can regulate cardiac dynamics, we next investigated the underlying neuronal substrate by testing whether a12 neuronal firing rates are related to spontaneous fluctuations in heart rate. We recorded from a12 neurons in two monkeys during resting state in the absence of any behavioral task (n=108 neurons; Methods). Consistent with previous work in this species (*41*), we observed that heart rate spontaneously fluctuates during rest on the timescale of seconds and minutes, generally between 2-3.5 Hz (**Fig. 2a**). In parallel, we observed that a substantial population of a12 neurons have firing rates that fluctuate over time with a striking resemblance to the heart rate. These spike-heart relationships could be either positive (**Fig. 2a**) or negative like a mirror reflection (**Fig. S6a**).

Importantly, this a12 activity is related to slow fluctuations in heart rate over time. It is not a simple sensory or motor signal encoding individual heartbeats. This could be seen by analyzing the power spectrum of spike-heart cross-correlations (**Fig. 2b**). The heartbeats themselves had a power spectrum highly concentrated on a 2-4 Hz frequency band matching the beat-to-beat rhythm of the heart (**Fig. 2b**, right, heart autocorrelation). They also had some power at lower frequencies, reflecting the slow fluctuations we observed in heart rate over seconds and minutes. If neurons encode heartbeats on a beat-by-beat basis, the spike-heart cross-correlation would have power above chance in the same 2-4 Hz band. Instead, the spike-heart cross-correlation was near chance in that band and only had significant power in a lower frequency band (**Fig. 2b**, 0.3-2 Hz). This suggests that a12 activity is related to slow changes in heart rate on the timescale of seconds or minutes.

To quantify this relationship, we modeled each neuron’s spiking probability at each millisecond of resting state as a function of its heart rate and other potential covariates (**Fig. 2c**). In particular, we included parameters in the model for heart rate estimated over the timescale of seconds (σ = 2 s) and minutes (σ = 60 s) (**Fig. 2c**, black curves; Methods, **Fig. S6**). To test whether seemingly heart rate-related activity might be better explained by other factors as arousal state, we also included analogous parameters for eye open/closed state (**Fig. 2c**, blue curves), which is a measure of arousal during resting state that predicts brain-wide functional connectivity patterns (*42*). To detrend the data and control for any potential effects of unrelated slow drifts in neuronal or heart signals, we also included covariates for linear, quadratic, and cubic effects of time (**Fig. 2c**, gray curves). To test each model and parameter for significance, we compared fits to the real data against fits to permuted datasets that preserved the detailed temporal structure of both spiking and heartbeat data but destroyed their relationship with each other, either by time-shifting the heart data by a random temporal offset (*43*) or by swapping heart data across sessions which produced the same pattern of results (**Fig. S6d-f**) (*44*, *45*).

We found that 27% of a12 neurons were best fit by including heart rate terms in the model (**Fig. 2d, left**). This was overwhelmingly driven by the term representing changes on the timescale of seconds, not minutes (**Fig. 2d, right; Fig. S6b,f**). The data was best fit by a timescale of 2-2.5 seconds (**Fig. 2e**). In addition, given our finding that a12 stimulation reduces heart rate, we hypothesized that a12 neurons with activity negatively related to heart rate may change their activity in advance of changes in heart rate. Indeed, we found that 13% of a12 neurons changed their activity more prominently before than after heartbeats while the opposite temporal pattern occurred at chance levels (**Fig. 2f; Fig. S7**), and these neurons were most often negatively related to heart rate (**Fig. 2g**). Thus, both stimulation and neuronal activity provide converging evidence for a functional role of a12 neurons in regulating heart rate.

### a12 and LHA can regulate heart rate on distinct timescales

Having established a12 and the LHA as prominent cardiac regulators in the primate brain, we next investigated the latency and state dependence of their influences on cardiac function. Our hypothesis is that LHA serves as a relatively fast, reliable, state-independent effector-controller, whereas a12 provides state-dependent top-down control of cardiac rhythm through the LHA, linking its cognitive function with cardiac output. To test this, we compared three key features of the stimulation-evoked heart rate response between a12 and LHA: the latency to response, the stimulation intensity threshold needed to evoke a significant response, and whether those responses were state-dependent or independent.

First, we stimulated the effective sites within a12 or LHA and reduced the current intensity strength to identify the threshold needed to evoke significant responses in either region. Stimulation at −300, −200, and −100 μA evoked responses in both areas, but stimulation at −50 μA evoked heart rate decrease in LHA but not a12, demonstrating that LHA requires a lower current intensity to evoke significant heart rate response compared to a12 (**Fig. 3a**; see **Fig. S8** for histograms). Next, we compared the latency of the effective-site heart rate response in response to electrical stimulation in either a12 or LHA. The response latency of LHA stimulation was significantly faster than that of a12 stimulation, indicating that LHA is more tightly linked to cardiac control (**Fig. 3b**). Finally, we evaluated the state dependency of the stimulation evoked responses in a12 and LHA by comparing the magnitude of the heart rate changes when the monkey had a low or high arousal state (see Methods). During a12 stimulation, when the monkey was in a high arousal state, the heart rate change was significantly attenuated (**Fig. 3c**). In contrast, during LHA stimulation, the monkey’s arousal state had no significant effect on the heart rate change. While activation of a12 and LHA both show very similar effects of the monkey’s heart function, these differences are consistent with the notion that LHA is downstream of a12 and anatomically closer to the heart. Altogether, these data show that the LHA serves as a fast, reliable, relatively state-independent effector-controller, whereas a12 provides a relatively more state-dependent anatomically top-down control of cardiac rhythm via LHA.

**Figure 3:**
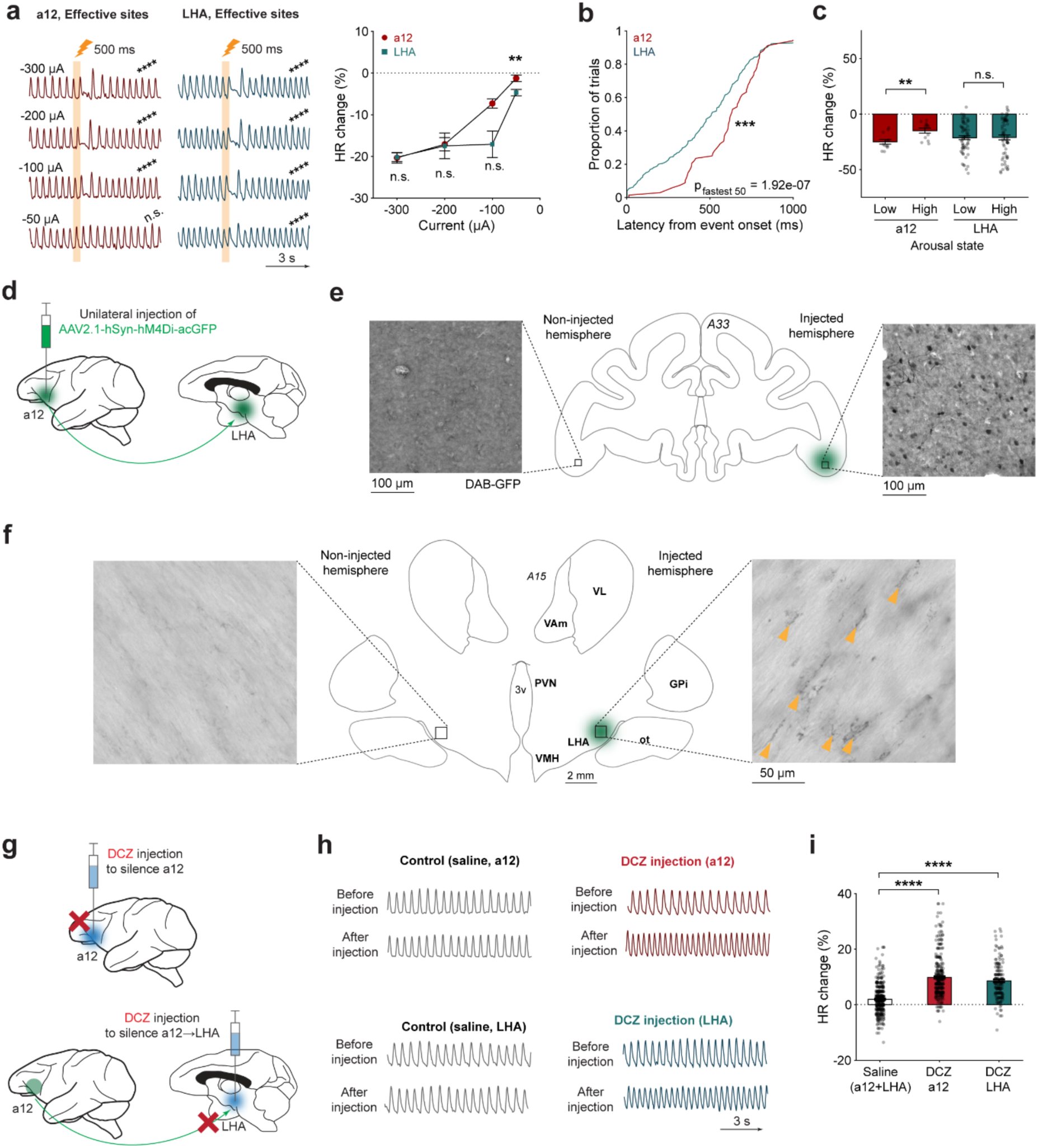
Dissecting the role of a12→LHA to modulate cardiac output. a, Left: Representative PPG traces as a function of current in the effective sites of a12 (red) and LHA (green). Conventions are the same as in Figure 1. Asterisks indicate significant decrease in the heart rate for each site averaged 0-1.5s after stimulation onset. For a12, n = 72, 46, 22, 48 stimulations for −300, −200, −100, −50 μA, respectively; P = 7.11e-13, P = 4.9e-09, P = 4.01e-05, P = 0.112. For LHA, n = 232, 29, 30, 30 stimulations; P = 4e-36, 3.17e-06, 2.13e-06, 1.8e-05 (two tailed signed-rank test against a hypothetical median of 0). Right: Average heart rate responses from a12 (red) and LHA (green) as a function of current. Data are mean ± s.e.m. P = 0.449, 0.443, 0.0507, 0.00176 for −300, −200, −100, −50 μA (two-tailed rank-sum test; a12 vs. LHA at each current). **b,** Cumulative distribution function plot of electrical stimulation-evoked heart rate response latencies in a12 (red) and LHA (green). n = 72, 232 for a12 and LHA stimulations, respectively. Asterisks indicates p-value when comparing all stimulations from a12 and LHA stimulation, while P_fastest 50_ indicates p-value when comparing stimulation response among the fastest 50% of stimulations from a12 and LHA. P = 0.000735; P_fastest 50_ = 1.92e-07 (two-tailed rank-sum test). **c,** Average heart rate changes in a12 and LHA evoked by electrical stimulation as a function of low or high arousal state. Data are mean ± s.e.m. For a12, n = 12 stimulations each for low and high arousal trials; P = 0.00355. For LHA, n = 64; P = 0.691 (two-tailed rank-sum test). **d,** Schematic illustration of the virus injection into a12, which will express hM4Di into both a12 cell bodies as well as a12 axon terminals in LHA. **e,** Representative histological images showing GFP expression in a12 cell bodies within the injected hemisphere compared to the non-injected hemisphere. **f,** Histology image showing GFP expression in LHA axon terminals projecting from a12 within the injected hemisphere compared to the non-injected hemisphere. Yellow arrows correspond to sites of labeled axon boutons. **g,** Schematic illustration of local DCZ injection to dampen neuronal activity. Top: DCZ injection into a12 to inactivate a12 neurons. Bottom: DCZ injection into LHA to dampen a12→LHA pathway. **h,** Top: Representative PPG traces before and after injection of saline or DCZ into a12. Bottom: PPG traces before and after injection of saline or DCZ into LHA. **i,** Average heart rate change after DREADD inactivation of a12 neurons and dampening of a12→LHA pathway. The heart rate changes after saline injection were pooled for both a12 and LHA. Data are mean ± s.e.m. (error bars too small to see); n = 399, 319, 161 measurements for saline, DCZ in a12, and DCZ in LHA. For saline vs DCZ in a12, P = 9.61e-53; for saline vs DCZ in LHA, P = 3.98e-28 (two-tailed rank-sum test). Throughout this figure, *, **, ***, **** indicate P < 0.05, 0.01, 0.001, 0.0001, respectively; n.s., non-significant.

Through electrical stimulation and neural activity analysis, we identified a prefrontal-hypothalamic system that exerts strong control over cardiac function in monkeys. Next, we asked if these neurons that specifically project from a12 to LHA are causally responsible for cardiac function. More specifically, what is the effect of inactivating these projection neurons on heart dynamics? To test this, we used a chemogenetic inhibition approach to specifically silence the axon terminals projecting from a12 and measured the resulting heart rate changes.

### Chemogenetic inactivation of a12→LHA modulates cardiac output

To evaluate the effect of a12→LHA pathway dampening on cardiac function in non-human primates, we employed a validated chemogenetic inactivation strategy (*46–48*). We injected viral vectors (AAV2.1-hSyn) encoding an inhibitory hM4Di DREADD (designer receptor exclusively activated by designer drugs) into a12 (**Fig. 3d**). This virus in particular is made for use in primates and is associated with DREADD-expression on both cell bodies as well as axon terminals (*46–48*). The expression of the DREADD at the site of the axon terminals in particular has previously allowed us and others to dampen the impact of the “injected brain region” on other brain regions by injecting the DREADD activator molecule deschloroclozapine (DCZ) into sites of the axon terminals expressing the hM4Di.

Histological examination of hM4Di-reporter GFP signals showed localized, dense expression within a12 cell bodies (**Fig. 3e**) as well as expression on LHA axon terminals in the injected hemisphere (**Fig. 3f**). These signals were not found in other control areas within the same plane as a12 or LHA (**Fig. S9**). Also, consistent with the anatomy in Figure 1, the location of GFP-labeled a12 axon terminals in LHA overlapped with the sites that displayed strong electrical stimulation evoked heart rate changes (**Fig. 1**).

After sufficient viral expression (> 1 month, Methods), we injected deschloroclozapine (DCZ) (*46*, *47*) into LHA (specifically the LHA regions where the a12 axon terminals were observed) and recorded the monkey’s heart rate change (**Fig. 3g**). As an additional control, we also injected DCZ into a12 to silence hM4Di-expressing a12 neurons independent of whether they projected to LHA or not.

To compute the effect of DCZ injection on cardiac dynamics, we calculated the heart rate change post-injection relative to the average heart rate pre-injection (**Fig. S10a**). While saline injections in either a12 or LHA produced no significant effects, DCZ injections in a12 and LHA both produced significant heart rate increases (**Fig. 3h**). On average, infusion of DCZ into a12 resulted in a heart rate increase of +9.82 ± 0.02%, while DCZ infusion into the area of LHA where a12 hM4Di expressing neurons were observed resulted in a heart rate increase of +8.56 ± 0.10%. Both of these effects were significantly greater than the saline controls within a12 and LHA pooled together (**Fig. 3i**; see **Fig. S10b-c** for individual animals). Together, these findings provide strong evidence that a12 controls heart rate in non-human primates and suggest it can do this through the a12→LHA pathway.

Area 12 is also strongly implicated in cognitive processes related to uncertainty and risk (*36*–*39*), and these cognitive features are tightly coupled to autonomic regulation and prominently associated with behavioral disruptions in mood disorders (*49–51*). Inspired by these observations, we next asked whether activity within a12 might serve as a site where cognitive and cardiac control intersect and interact. We aimed to address this question using techniques that may be later translated for the clinic. To address these needs and questions, we added to our existing toolkit of causal manipulation using DREADDS and further developed a sonogenetic approach that enables genetically targeted, temporally restricted, and relatively noninvasive perturbation of area 12 that may have the potential to translate a12 neuromodulation towards future development of clinical treatment paradigms. We then used this method to causally modulate heart rate (**Fig. 4**) and test the impact of a12 perturbations on decision-making processes (**Fig. 5**). Finally, we probed the link between a12 encoding of decision variables and of moment-by-moment fluctuations in cardiac state (**Fig. 5**).

**Figure 4:**
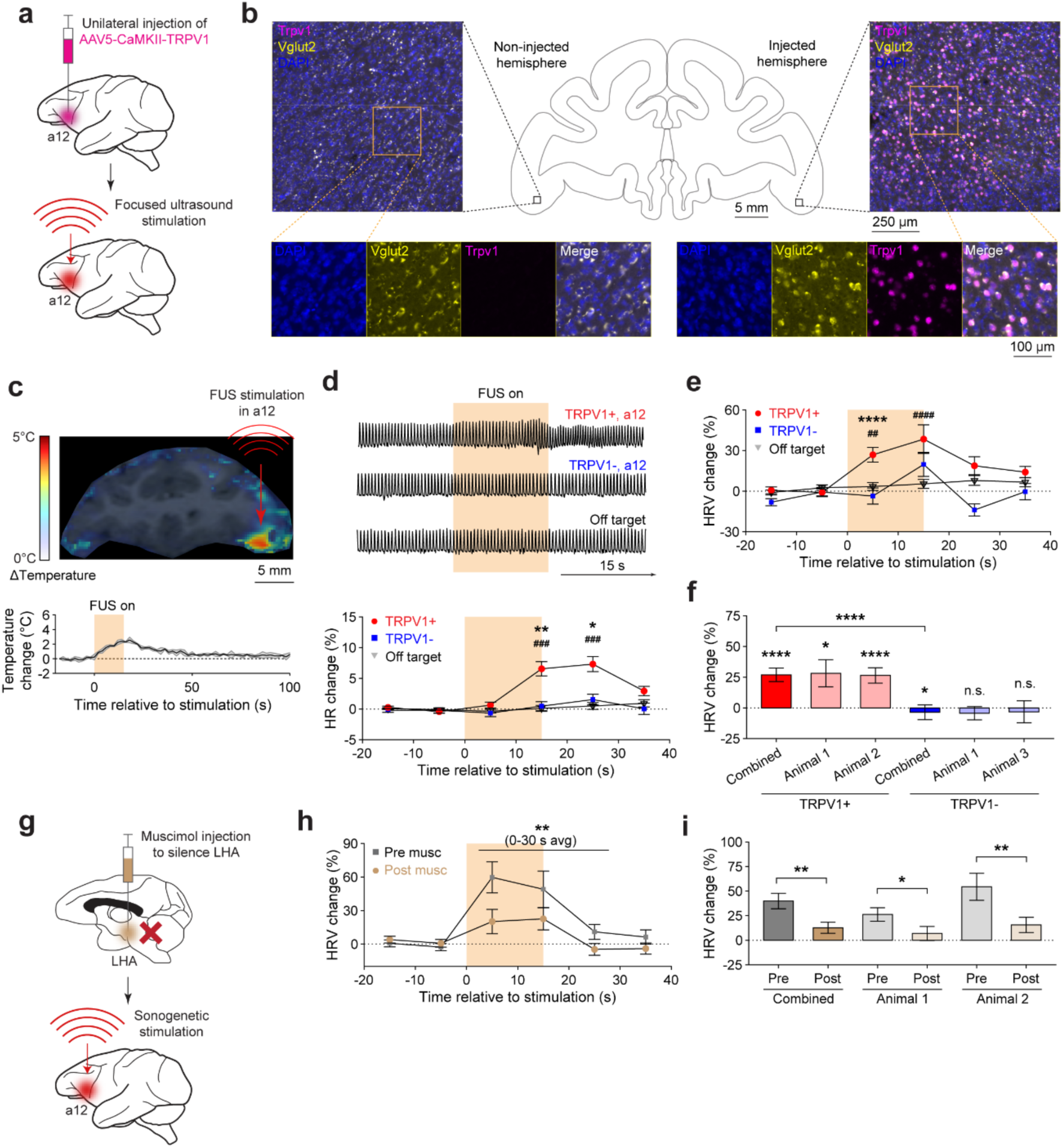
Sonogenetic perturbation of a12 modulates heart function. a, Schematic illustration of the sonogenetic experimental procedure performed in non-human primates, which first involves an injection of AAV encoding TRPV1 into a12 followed by focused ultrasound (FUS) stimulating the same brain region. **b,** Representative fluorescent *in situ* hybridization images in a12 visualizing DAPI (blue), Vglut2 (yellow), and Trpv1 (magenta) expression in both the injected and non-injected hemispheres. **c,** Top: representative MR thermometry map showing localized heating in a12. Bottom: Temperature change time course computed with an ROI within a12. The yellow bar indicates FUS stimulation period. Shaded region represents s.e.m. **d,** Top: Representative PPG traces shown for a12 TRPV1+ (red), a12 TRPV1-(blue), and off-target (dorsal prefrontal cortex, grey) groups. Bottom: Time course of the average heart rate change computed in 10 s time bins for the same three groups. Yellow bar indicates FUS stimulation period. Data are mean ± s.e.m. * denotes comparison between TRPV1+ and TRPV1-groups. # denotes comparison between TRPV1+ and off-target groups. n = 87, 55, 135 stimulations for TRPV1+, TRPV1-, and off-target. For 0-10 s time window, P = 0.167, 0.146 for TRPV1+ vs TRPV1-and TRPV1+ vs off-target, respectively; for 10-20 s time window, P = 0.000911, 1.09e-05 (two-tailed rank-sum test). **e,** Time course of the average heart rate variability changes computed in 10 s time bins for the same three groups. Yellow bar indicates FUS stimulation period. Data are mean ± s.e.m. For 0-10 s time window, P = 4.61e-05, 0.000573 for TRPV1+ vs TRPV1-and TRPV1+ vs off-target, respectively; for 10-20 s time window, P = 0.325, 0.00669 (two-tailed rank-sum test). **f,** Average heart rate variability changes 0-10s after stimulation onset, comparing TRPV1+ and TRPV1-responses on a subject-by-subject basis. Data are mean ± s.e.m. Asterisks above each bar indicate a signed-rank test against a hypothetical median of zero. n = 21, 66, 20, 35 stimulations for animal 1 (TRPV1+), animal 2 (TRPV1+), animal 1 (TRPV1-), animal 3 (TRPV1-), respectively. In the TRPV1+ group, P = 4.59e-05, 0.0457, 0.00036 for combined animals, animal 1, animal 2, respectively; In the TRPV1-group, P = 0.027, 0.37, 0.0533 for combined animals, animal 1, animal 3. Asterisks above the bracket indicate rank-sum test. P = 4.61e-05. **g,** Schematic illustration of the LHA inactivation experiment during sonogenetic stimulation of a12. **h,** Time course of the average heart rate variability change computed in 10s time bins for the same groups. The yellow bar indicates FUS stimulation period. Data are mean ± s.e.m. Asterisks above the line indicate rank-sum test computed between pre-and post-muscimol responses averaged over 0-30 s in three consecutive 10 s time bins. n = 37, 54 for pre-and post muscimol stimulations; P = 0.0031 (two-tailed rank-sum test). **i,** Average heart rate variability changes 0-30s after stimulation onset, comparing sonogenetic-evoked responses pre-and post-muscimol injection on a subject-by-subject basis. Data are mean ± s.e.m. Asterisks above the bracket indicate rank-sum test. n = 19, 18, 18, 36 stimulations for pre-muscimol (animal 1), post-muscimol (animal 1), pre-muscimol (animal 2), post-muscimol (animal 2). P = 0.0347, 0.0089 for animal 1 and animal 2 (two-tailed rank-sum test). Throughout this figure, *, **, ***, **** indicate P < 0.05, 0.01, 0.001, 0.0001, respectively; n.s., non-significant.

**Figure 5:**
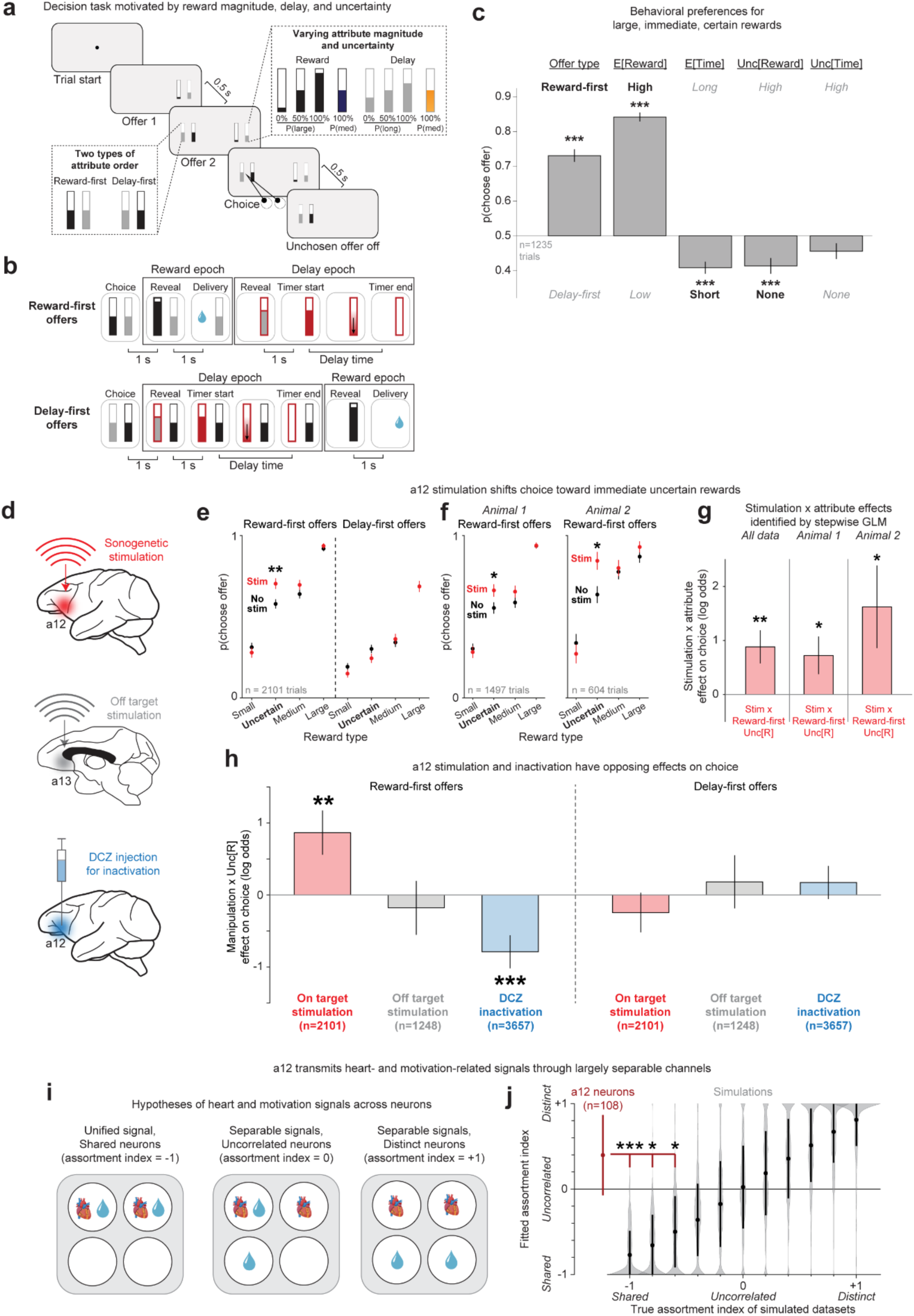
Sonogenetic and chemogenetic perturbation of a12 bidirectionally modulate motivated behavior. **a**, Decision task motivated by reward magnitude, delay, and uncertainty. Animals chose between offers which provided different reward magnitudes, delay durations, and orders of reward and delay (reward-first or delay-first). Both rewards and delays could be either certain or uncertain. **b**, Sequence of events after choosing reward-first vs. delay-first offers. **c**, Animals were highly motivated to perform the task, shown by frequently choosing offers that were reward-first, high E[R], low E[T], and low Unc[R]. *** indicates p < 0.001 (two-tailed signed-rank test). **d**, In different sessions, animals performed the task during sonogenetic on-target stimulation of a12 (red), off-target stimulation of a13 (gray), or DCZ to silence a12 (blue). **e**, On-target stimulation of a12 shifts choice toward uncertain immediate rewards. Probability of choosing an offer as a function of reward magnitude (x-axis: small, uncertain, medium, or large) and attribute order (reward-first, left; delay-first, right), separately for stimulation trials (red) and control trials (black). ** indicates significant difference (p < 0.01, rank-sum test). **f**, This result occurred in both animals (left vs. right; same format, showing data for reward-first offers). **g**, Consistent results from stepwise GLM, which identified only one Stimulation x Attribute interaction effect (Stim x Reward-first Unc[R]). This occurred consistently in pooled data (left) and each individual animal (middle, right). *, ** indicates significant differences at p < 0.05, 0.01 (two-tailed t-test). **h**, a12 bidirectionally regulates choice, indicated by Manipulation x Reward-first Unc[R] effect (left) being positive for on-target stimulation (red, same as above), absent for off-target stimulation (gray), and negative for DCZ inactivation (blue). As a control, no similar results occurred for a potential Manipulation x Delay-first Unc[R] effect (right). **, *** indicates p < 0.01, 0.001 (two-tailed t-test). **i**, Hypotheses for a12 transmitting heart-and motivation-related signals as a single unified signal via a shared neuronal substrate (left), or separably via uncorrelated (middle) or distinct (right) substrates, corresponding to assortment indexes of −1, 0, or +1. **j**, Fitted assortment index for a12 neuronal population (red, error bar indicates +/− 1 bootstrap SE) and simulated datasets generated with different known assortment indexes spanning its full possible range (black; error bar indicates +/− 1 SD; gray violin plots show all individual simulations, n=400 per data point). a12 neurons had largely separable signals, with a significantly higher assortment index than would be expected based on simulations with a true index of −1, −0.8, or −0.6. *, *** indicates p < 0.05, 0.001 (two-tailed tests comparing real vs. simulation distribution).

### Sonogenetic perturbation of a12 modulates heart rate dynamics

To test whether a less invasive perturbation of area 12 can modulate cardiac dynamics and cognitive state in primates, we sought a method that combines genetic targeting with the ability to perturb relatively large volumes of deep cortex in a temporally restricted manner. Genetically based neuromodulation tools, such as optogenetics and chemogenetics, have provided powerful means for cell-type-specific circuit perturbation. In primates, however, optogenetic manipulation of deep prefrontal regions typically requires implantation of optical hardware. Moreover, the robust behavioral and physiological effects thought to be necessary for clinical impact often require perturbing relatively large tissue volumes, which raises further challenges of light penetration and volume coverage (*52*). Chemogenetics is comparatively less invasive for systemic applications, but has limited temporal resolution due to the long residence time of the activating ligand, which could be appropriate for many use cases that require sustained modulation (*53*), but also could be longer than desired for some other clinical applications.

Focused ultrasound (FUS) has emerged as a method for noninvasively perturbing neural activity over large spatial scales, although the mechanisms engaged by FUS remain under investigation (*54–56*). Modulating heart rate dynamics and internal states is a key direction for clinical neural modulation development. Guided by prior findings, we combined FUS with molecular techniques and tested whether it can regulate heart rate dynamics through a12. This approach, termed sonogenetics, uses virally expressed ultrasound-sensitive actuators within a neuron population of interest to increase cell-type specificity while retaining the ability to modulate relatively large and deep cortical volumes (*57*, *58*).

Since sonogenetics remains at an early stage of development with only a limited number of candidate actuators available (*59–62*), we used the thermosensitive ion channel TRPV1 as a key initial test of feasibility because TRPV1-based sonogenetics has previously produced robust neuronal activation and behavioral modulation in rodents (*63–65*). This approach relies on mild, spatially confined FUS-induced tissue warmed to activate TRPV1. Here, our objective was not to establish TRPV1 as the final clinical actuator, but to determine whether sonogenetic modulation of a12 is feasible in non-human primates and can influence heart rate dynamics.

Specifically, we developed a TRPV1-based sonogenetic approach in which neurons express the thermosensitive ion channel TRPV1, enabling selective perturbation by mild, spatially confined ultrasound-induced mild warming. We unilaterally injected viral vectors into area 12 to express TRPV1 in mostly excitatory neurons, and after ∼1-2 months, applied FUS targeted to the injection site and studied heart rate responses to this manipulation (**Fig. 4a**).

We verified the expression of exogenous TRPV1 in the monkey brain and assessed the mild warming resulting from FUS stimulating a12. Using fluorescent in situ hybridization, we confirmed that exogenous Trpv1 mRNA was expressed within a12, was spatially confined to a12, and was mostly co-expressed with Vglut2, a gene marker present in excitatory neuron populations (**Fig. 4b**). We visualized the overlap between TRPV1 and Vglut2 (**Fig. S11a-b**), which showed that our CaMKII-promoter strategy to target excitatory neurons within cortical regions was approximately 82 ± 3% and was relatively on par with prior literature (*66*, *67*). Furthermore, we did not observe Trpv1 mRNA in the non-injected hemisphere or other regions within the prefrontal cortex (**Fig. S11c**). To target a12 with FUS, we initially chose stimulation parameters that resulted in mild warming to activate TRPV1, based on previous sonogenetic studies conducted in rodents for neuronal and behavior manipulation (**Fig. S12a**). We then confirmed the targeting of FUS and the degree of FUS-related warming by a standard non-invasive technique used in neurosurgical settings called magnetic resonance (MR) thermometry (*68*). The MR temperature map shows mild warming within area 12, reaching an estimated peak temperature increase of 2.5°C (**Fig. 4c**). To avoid the possibility of overheating the brain due to consecutive stimulations, we set the inter-stimulation interval to be in the range of 100 to 185 s to ensure that the brain temperature returned to baseline before performing another stimulation. This stimulation protocol did not produce any changes in edema or compromised brain structures assayed by T2-weighted MR scans. We point out that simple warming of neurons and the effects of FUS alone may change neural activity (*55*, *69*); therefore, we performed the stimulation in three groups and compared their heart rate responses: stimulation in a12 with TRPV1 expression (TRPV1+), stimulation in a12 without TRPV1 expression (TRPV1-), and stimulation in a dorsal prefrontal area without TRPV1 expression (off-target).

To evaluate the heart rate responses, we computed the time course of the mean heart rate and heart rate variability changes for each group, quantified over 10 s time bins (**Fig. 4d-e**). The most consistent effect of sonogenetic stimulation was on heart rate variability. In the TRPV1+ group, heart rate variability increased significantly in both animals, and this increase occurred within the first 10 s after stimulation onset — that is, during the 15 s stimulation itself (**Fig. 4d-f**, **Fig. S12**). This is consistent with the large increases in heart rate variability that also accompanied electrical stimulation of a12 and LHA (**Fig. 1f**, **Fig. S3f**). In rodents, TRPV1-mediated sonogenetic activation delivered with similar ultrasound parameters has an onset on the order of a few seconds, with neural responses beginning shortly after stimulation onset and behavioral effects occurring during the stimulation period (*63*, *65*), which aligns well with our results here. This is a notably more sluggish, and likely more variable, timescale than that of electrical stimulation, plausibly due to the gradual warming we measured by MR thermometry (**Fig. 4c**). In contrast, the TRPV1-group yielded substantially smaller effects (**Fig. 4d-f**). There was a slight increase in heart rate variability 10-20 s after stimulation onset, and there was no significant difference in heart rate variability between TRPV1+ and TRPV1-groups in this later time bin (**Fig. 4e**). This suggests that FUS stimulation without TRPV1 expression could also affect the heart through a12, albeit with weaker magnitude and later onset than those observed from the TRPV1+ group.

In addition, sonogenetic stimulation significantly increased heart rate in one of the two TRPV1+ animals (**Fig. 4d**, **Fig. S12b-c**), and this substantial increase emerged after a delay, approximately 10-30 s after stimulation onset (**Fig. 4d**). Because the stimulation lasted 15 s, this window begins near the end of the stimulus and extends well beyond its offset. The delayed heart rate increase could reflect a rebound following the initial autonomic response, analogous to the positive heart rate excursion in the late window after electrical stimulation of a12 and LHA (**Fig. 1d,m**), though that late electrical effect did not itself reach significance relative to ineffective site stimulation. Alternatively, since FUS-induced warming decayed over tens of seconds after the stimulation ended (**Fig. 4c**), continued TRPV1 activation beyond the stimulation period and potential thermo-sensing by endogenous processes cannot be excluded.

To investigate whether LHA plays a role during sonogenetic stimulation of a12, we administered muscimol to the subregions within LHA that we previously found regulated cardiac function and tested whether this altered sonogenetic-evoked heart rate responses (**Fig. 4g**). LHA inhibition increased the overall heart rate (**Fig. 1h**) but also significantly attenuated the magnitudes of sonogenetic-evoked changes in heart rate variability (**Fig. 4h-i**). The reduction in a12-evoked responses, due to a state change mediated by LHA inhibition, may be related to the state-dependent nature of a12’s influence on cardiac activity, analogous to the state-dependent responses we observed from electrical stimulation within the same brain region (**Fig. 3c**). Overall, these findings suggest that LHA plays a role in a12 mediated heart rate changes evoked by TRPV1-mediated sonogenetics.

### Sonogenetic and chemogenetic perturbation of a12 bidirectionally modulate motivated behavior

In addition to regulating the heart, a12 has also been implicated in regulating motivated behavior (*36–39*). We therefore asked whether the sonogenetic stimulation that we demonstrated influences the heart might also influence motivated behavior. To do this, we trained two monkeys to perform a task allowing them to choose between juice rewards with different magnitudes, timing, and uncertainty (**Fig. 5a**). On each trial the animal chose between two options. Each option consisted of two bars with different colors. One bar indicated the possible reward magnitudes (*R*). The reward could be either *certain*, with a single specified magnitude (small, medium, or large) or *uncertain,* with two possible magnitudes (50/50 chance of small or large). The other bar indicated the possible time delay durations (*T*), which in an exactly analogous manner, could be certain (short, medium, or long) or uncertain (50/50 chance of short or long). Finally, the order of the reward and delay bars on the screen indicated the order the reward and delay would be delivered (**Fig. 5b**). Reward-first offers delivered the reward and then the delay, while delay-first offers delivered the delay and then the reward. As expected, monkeys showed strong conventional preferences for large immediate rewards (**Fig. 5c**; similar in each animal, **Fig. S13a**). They strongly preferred offers with high expected reward (high ≻ low E[R]), short expected delays (short ≻ long E[T]), and where the reward was delivered before the delay (Reward-first ≻ Delay-first). They also tended to avoid uncertain rewards (low > high Unc[R]), which has been previously observed in this species in some tasks (*70*) though not others (*71*).

We found that a12 sonogenetic stimulation shifted choices toward immediate uncertain rewards. Stimulation was delivered on a trial-by-trial basis (**Fig. 5d**, schematic highlighted in red; see Methods for parameters and stimulation timing). On trials with stimulation, animals were significantly more likely to choose reward-first uncertain reward offers (**Fig. 5e**). This result occurred consistently in both animals (**Fig. 5f**). Furthermore, this result did not occur for certain rewards of any magnitude (small, medium, or large), nor did it occur for delay-first offers (**Fig. 5e**, **Fig. S13b**). We also found similar results using a formal modeling approach to account for all motivational attributes and stimulation effects simultaneously (**Fig. 5g**). We modeled each animal’s behavior using a generalized linear model including terms quantifying the main effect of a12 stimulation and of nine distinct offer attributes: offer type (reward-first or delay-first), and E[R], E[T], Unc[R], Unc[T] separately for reward-first and delay-first offers. We then used stepwise regression to identify the set of stimulation x attribute interaction terms that significantly improved the fit. This procedure identified a single significant interaction term: a stimulation x Unc[R] effect for reward-first offers. The same result occurred regardless of whether this procedure was applied to the entire dataset or to each individual animal (**Fig. 5g**). This effect was present when analyzing the data in either a pooled manner or on a session-by-session basis (**Fig. S13c**).

Given that a12 sonogenetic stimulation and DCZ inactivation may have partially opposing effects on the heart, we asked whether they might also have opposing effects on motivated decisions (**Fig. 5d**, schematic highlighted in blue). Furthermore, if sonogenetic stimulation has its effect through the intended mechanism of activating TRPV1 channels in the targeted brain region (a12), it should not have any effect when delivered to a different, off-target region of prefrontal cortex (a13; **Fig. 5d**, schematic highlighted in gray). To test this, we conducted additional experimental sessions using either off target sonogenetic stimulation or DCZ inactivation. We analyzed behavior using the same model identified above, in which manipulation effects were represented by the identified interaction term (“manipulation x Unc[R] for reward-first offers”; **Fig. 5h**, left). As a further control, we included an additional, analogous interaction term for delay-first offers (“manipulation x Unc[R] for delay-first offers”; **Fig. 5h**, right). We found that sonogenetic stimulation and DCZ inactivation had opposing effects on choice of reward-first uncertain offers. Choice was significantly increased by stimulation, not significantly affected by off-target stimulation, and significantly reduced by DCZ inactivation (**Fig. 5h**, left). There were no similar effects for delay-first offers (**Fig. 5h**, right). Stimulation and inactivation effects were significantly larger on reward-first offers than delay-first offers (stimulation: p = 0.0056; DCZ: p = 0.0019; off-target: p = 0.62; *t*-tests). Similar results could be seen in raw choice percentages, and in an extended GLM including all possible manipulation x attribute effects, which both similarly indicated that stimulation and inactivation had opposing effects on choice (**Fig. S13d**).

Finally, given our findings that a12 regulates both the heart and motivated decisions, we asked whether their underlying neuronal substrates might be conjoined with each other. In particular, we wanted to know whether a12 transmits heart-and motivation-related activity using a single unified signal via a shared subpopulation of neurons, which would be suitable to regulate the heart and motivation in a specific, stereotyped relationship with each other (**Fig. 5i**, “Unified signal, Shared neurons”). Alternately, a12 could transmit separable signals, which would be suitable to regulate the heart and motivation in a flexible manner, adjusting them differently depending on the context (**Fig. 5i**, “Separable signals”). This could be accomplished by several possible neuronal substrates, such as by partially overlapping subpopulations of neurons where the presence of one signal is uncorrelated with the other (**Fig. 5i**, “Uncorrelated neurons”), or by a labeled-line code with distinct, non-overlapping subpopulations of neurons for heart vs. motivation (**Fig. 5i**, “Distinct neurons”).

To test this, we recorded the activity of the same n=108 a12 neurons while monkeys performed the task and quantified their motivation-related activity preceding the choice and its significance using a standard generalized linear model (Methods). We then used a modeling approach to estimate an *assortment index* which quantifies the separability of a12 heart-and motivation-related activity, while controlling for the base rates of each type of activity, and for the rates of statistical false positives and false negatives that may arise from each neuron’s sample size and sampling noise (Methods). We defined the assortment index as +1 if the presence of heart-and motivation-related activity are maximally separable from each other, 0 if they are uncorrelated with each other, and −1 if they are minimally separable from each other (**Fig. 5i**). We confirmed that this model recovered indexes that closely resembled the true indexes of simulated datasets with closely matched statistical structure to the real data, over a wide range of possible assortment levels (**Fig. 5j**, gray). When we then applied the same model to the real data, it recovered an assortment index of +0.4, consistent with separable signals midway between the “Uncorrelated” and “Distinct” hypotheses (**Fig. 5j**, red). This was significantly greater than all possible indexes in the vicinity of the “Shared” hypothesis, that is, indexes below −0.5 (**Fig. 5j**). This lack of correlation between signals was highly robust. It occurred for activity in all task epochs, and for activity related to each motivational variable within each epoch (**Fig. S14**). Thus, our data suggests that a12 contains neurons with both heart-and motivation-related signals and uses a neural code making it possible to regulate those functions separably.

## DISCUSSION

Our heart rate is linked to many different cognitive and emotional states, yet the underlying mechanisms and neural substrates governing the flexible coordination between the heart and cognition remain incompletely understood. Here, we identify a pathway by which a12, a region of primate prefrontal cortex that regulates cognition and decision making, also governs heart function through its projection to LHA. Recent studies provide evidence that a12 is an important region for behavioral control under uncertainty, with particular contributions to decision making among probabilistic options (*37*, *38*), credit assignment (*39*), and information seeking to resolve uncertainty (*36*). Uncertainty plays a key role in the dysregulation of cognitive-and mood-disorders (*49–51*), which are also associated with dysregulation of autonomic and heart dynamics (*40*, *72*, *73*). Our findings show that a12 regulates the dynamics of the heart, providing a potential substrate for the flexible coordination of cognition and heart control within the primate prefrontal cortex. Neurons in a12 closely tracked moment-to-moment heart rate fluctuations (**Fig. 2**), and their causal perturbation by sonogenetics and chemogenetics impacted risky decision making under uncertainty (**Fig. 5**).

a12 controls heart rate at least in strong part through its inputs to LHA. It is well established that the LHA is responsible for arousal and autonomic regulation of visceral organs in rodents (*74*–*76*). Here, we show that the primate LHA also contains a relatively spatially restricted region that can robustly modulate heart rate (**Fig. 1**), that activation of this region can change heart rate without always changing all the other features of the arousal (**Fig. S4-S5**), and that dampening of a12 inputs to LHA changes heart rate (**Fig. 3**). Indeed, among prefrontal regions that project to LHA (*31–34*), causal manipulation of a12 produced the strongest effects on heart rate dynamics.

We also found that stimulation of more dorsal regions of the hypothalamus did not produce the same rapid suppression of heart rate that we found when stimulating LHA. However, these areas may still impact the autonomic system on a distinct, slower timescale through secretory and other pathways (*77*). Dorsal hypothalamic areas in primates receive inputs from the ventral medial prefrontal cortex (vmPFC), including areas 14 and 25 (*31*, *33*, *35*), that are known to regulate internal or emotional states over slow timescales (*78–83*). Therefore, how the rapid a12→LHA circuit and the dorsal hypothalamus circuits (potentially receiving vmPFC and other autonomic-related inputs (*84*, *85*)) interact to control heart rate requires further examination. Moreover, even within primate area 12, there are reported differences in how neurons process decision variables. Future work must examine the relationship of these functional gradients and heart rate regulation (*86*).

The heart rate effects evoked by a12 and LHA stimulation also differed in ways roughly consistent with a hierarchically arranged control system. LHA required a lower current intensity to evoke a heart rate response, responded with a shorter latency, and was insensitive to the animal’s arousal state, whereas a12 effects were detected later (they were slower), required stronger stimulation, and were attenuated when the arousal was relatively high (**Fig. 3**). Together, these data are consistent with LHA acting as a fast effector through which a12 exerts slower, more state-dependent control of the heart.

Mammalian insula is known to sense changes in heart-rate related dynamics and regulate internal states (*87–89*), but how remains poorly understood. In primates, a12 is located close to and is strongly interconnected with the anterior insular cortex (*90*), and prior work showed that very strong microstimulation of the anterior insula produced heart rate modulations along with consistent emotion-related facial and body responses (*91*). In comparison, our current study revealed that microstimulation in a12 or LHA evoked similar heart rate effects without producing consistent significant movement responses, and a12 neurons covaried with heart rate dynamics across states in a manner that could not be explained by arousal state alone. Furthermore, stimulation related heart rate effects occurred even when eyes were closed. These results imply that a12 and insula could engage related but distinct pathways to flexibly modulate autonomic activity depending on emotional state. To understand how heart rate modulation is coordinated with the outward expression of emotion observed during insular stimulation requires a new investigation of how a12→LHA interacts with the insular cortex at the anatomical, neurophysiological, and causal levels.

We found that a12 neurons encode heart-and motivation-related variables in largely separable manners (**Fig. 5**). This is what one would want for a neural hub that links motivation to autonomic control. a12 has the potential to separately monitor and regulate motivational and autonomic processes, while still retaining the ability to coordinate them with each other in a flexible, context-dependent manner (by communication between the subsets of motivation-related vs. heart-related neurons; or via the subset of neurons significantly related to both processes). This coordination could also occur through circuitry downstream of a12. For example, information seeking or uncertainty related neurons may project to the basal ganglia, through the a12→basal ganglia→lateral habenula (LHb) circuit (*92*, *93*), while heart rate sensitive neurons may project to the LHA. It has also been reported that LHb and LHA interact and relay motivation related signals (*94*, *95*), which could further allow autonomic and cognitive regulation to intersect at the level of the LHb↔LHA pathway.

We report the first study of TRPV1-sonogenetics to modulate primate brain circuits (**Fig. 4**). By transfecting CaMKII+ neurons in area 12 to express TRPV1 and applying mild warming to the region via FUS, we evoked early, large increases in heart rate variability that are much greater than what occurs from FUS stimulation without TRPV1 or in off-target controls. Notably, the increase in heart rate variability occurred during the ultrasound stimulation itself, whereas the substantial increase in heart rate in one animal emerged later (**Fig. 4d,e**) and may represent a rebound effect also observed in electrical stimulation. However, sonogenetic perturbation is different from electrical stimulation due to the spatial extent of the affected area, differences in neuronal populations being engaged, and the temporal resolution of these techniques (*58*, *60*, *61*, *96*) – with TRPV1-sonogenetics eliciting a slower and relatively more variable change in neural dynamics than the fast electrical stimulation or optogenetics (*63*, *65*). Potential timecourses and other differences are therefore expected to vary across electrical stimulation, optogenetics, and sonogenetics, and further understanding of circuit systems-level mechanisms for all three are required to fully dissociate them.

Additionally, FUS alone can modulate brain activity, and the precise parameters that are required for this remain under investigation (*54–56*, *97*). Typical FUS parameters largely differ from those used in the current study in that these parameters were designed to induce mild warming in the brain. Indeed, several mechanisms of FUS activation of neurons without expression of exogenous proteins have been proposed (mechanical activation of endogenous mechanosensors, thermal activation of endogenous thermosensors, and intramembrane cavitation). These mechanisms could contribute to autonomic modulation in diverse ways and are critical for understanding, interpreting, and optimizing sonogenetic effects for clinical applications. In our study, we used off-target and TRPV1-controls to highlight the unique contribution of TRPV1+ and also show that a12 can be a future clinically relevant target for treatment of disorders associated with heart and cognitive dysfunction, one that is potentially accessible enough to be a target for noninvasive transcranial approaches in humans.

The bi-directional connection between the brain and visceral organs remains a key area of research to uncover the neural mechanisms of autonomic regulation in mammals. The prefrontal-hypothalamic pathway we have characterized provides a neural substrate through which the cognitive and motivational systems that govern decision-making can interface with subcortical autonomic control systems to govern the heart. More broadly, identifying a defined, manipulable circuit for top-down cardiac control in the primate brain provides a foundation for understanding how cognitive and emotional states come to shape bodily physiology, and a potential target for therapies aimed at restoring healthy brain–heart coupling in psychiatric and cardiovascular disease.

## METHODS

### Experimental model and study participant details

#### Animal procedures

All animal procedures were performed under a protocol approved by the Washington University in St. Louis Institutional Animal Care and Use Committee (protocol number 22-0164). Three adult male rhesus monkeys (*Macaca mulatta*; animals 1, 2, and 3; aged 10–12 years old) were used for the experiments. Their experimental allocations are summarized in Table 1. This sample size is standard for behavioral and neurophysiological studies in non-human primates. A customized plastic head holder and plastic recording chamber were implanted on the skull with dental acrylic under general anesthesia and sterile surgical conditions. Craniotomy chambers were tilted and aimed at the prefrontal cortex and hypothalamus. After the monkeys recovered from surgery, they participated in behavioral, neurophysiological, and circuit manipulation experiments.

#### Heart rate acquisition

The monkey heart rate was measured using photoplethysmography (PPG), a noninvasive optical approach to measure heart rate based on blood volume changes in the vasculature. A PPG device (PulseSensor, World Famous Electronics) was attached to the monkey’s earlobe using an ear clip, which provides a signal with a tight relationship to the electrocardiogram and is well suited for use in awake, behaving macaques (*98–100*). The signals from the device were first configured to a microcontroller board (Arduino Uno Rev3), then relayed to a data acquisition system (Plexon).

#### Auxiliary behavior acquisition

In addition to heart rate, we acquired several other behavioral parameters from the monkey, such as eye tracking data and force transducer movement data to quantify overall body movement. Eye position and pupil size were acquired with an infrared eye-tracking system (Eyelink, SR Research). All these signals were relayed to the same acquisition system (Plexon) for synchronized data recording.

#### Electrical stimulation

Epoxy-coated microelectrodes (FHC, tungsten, cat no. UEWLFESEENNE) were used to deliver electrical current. These microelectrodes were modified to lower their impedances to a range within 100 and 200 kOhms. Stimulation sites were determined with a 1 mm spacing grid system and with the aid of anatomical magnetic resonance images. This MRI-based prediction of stimulation sites was facilitated by custom-built software (PyElectrode (*101*)) and histology. Electrodes were inserted into the brain through a stainless-steel guide tube and advanced by an oil-driven micromanipulator (MO-97A, Narishige). While lowering the electrode to the target stimulation site, the electrode was connected to the recording system (Plexon) to visualize neural signals and roughly confirm the MRI-predicted trajectory of the electrode track via electrophysiology. Within a12, Cg, a13; the stimulation sites were between interaural A∼32 to A∼35. Within the hypothalamus they were between A∼14-A∼17.

Once lowered to the target site, the electrode was then connected to a stimulator (A-M Systems, Model 4100 Stimulator). A TTL-pulse from a data acquisition box (DATAPixx) triggered the stimulator to deliver a stimulation train (400 μs total pulse duration across both phases, biphasic symmetric, cathodal-first; 400 Hz pulse frequency; total train duration of 500 ms; stimulation current varied between −300 and −50 μA, reported throughout as the amplitude of the leading cathodal phase; **Fig. S1a**) at a randomly selected inter-stimulation interval between 45 and 90 s. Heart rate and other auxiliary behaviors were continuously acquired during this stimulation protocol for downstream analyses.

#### Visualization of BDA anterograde tracer slides

In brief, 0.2 μL of biotinylated dextran amine (BDA, 10000 MW, Molecular Probes) was pressure injected into a12. After tissue sectioning, slices were processed directly with the avidin–biotin–peroxidase method (Vector ABC) with silver/gold intensification. Slides were then imaged and digitized using a typical brightfield microscope. This was cs. # om31 from the collection of Dr. Joel Price.

#### Injection of viral vectors

For chemogenetic manipulation of neurons, adeno-associated viral vectors (AAV) encoding the designer receptor exclusively activated by designer drug (DREADD) hM4Di were used (AAV2.1-hSynI-Flag-hM4Di-IRES2-AcGFP, 2e13 vg/mL). This AAV was provided by the Innovative Brain Virus Vector Core (Takada Group, Kyoto University, Japan). For sonogenetic manipulation of neurons, AAVs encoding TRPV1 (AAV5-CaMKII-TRPV1-DsRed, 1.2e12 vg/mL) were used. Cloning, packaging, purification, and viral titer calculations for this viral vector were performed by the Hope Center Viral Vectors Core at Washington University School of Medicine.

AAVs were unilaterally injected into a12 using a 10 uL microsyringe (Hamilton) combined with a custom-made injection needle. 2-4 injection sites were identified based on heart rate sensitivity assayed with electrical stimulation. For each site, 3-5 uL of AAVs were injected at a rate of 0.030 uL/min. Once the injection was complete, the injection needle stayed in the injection site for 1 hour to minimize upward leakage along the injectrode track and ensure AAV uptake in the brain.

#### Chemogenetic inactivation

Deschloroclozapine (DCZ) was used to inactivate hM4Di-expressing a12 neurons as well as a12 axon terminals within LHA. 2.5 mg DCZ (Bio-Techne) in 100 µL DMSO (Bio-Techne) stock solution was prepared and then diluted in 1X PBS for a final concentration of 100 nM DCZ on the day of use. This solution was unilaterally injected into either a12 or LHA of the virus-injected hemisphere. Injection sites were chosen based on heart rate sensitivity assayed with electrical stimulation. DCZ (3–5 uL) was infused using a 32-or 64-channel linear array with a fluid capillary (v-probe, Plexon) at a rate of 0.1-0.13 µL/min. Saline (3 µL) was injected in either a12 or LHA as control sessions. DCZ and control sessions were intermingled to prevent the animal from predicting the nature of the experiment. Experiments were also performed with a minimum interval of 4 days to ensure the DCZ was washed out. The pre-injection heart rate was recorded from the time the injectrode reached the target injection site until injection onset; the post-injection session started once the injection was completed (**Fig. S10a**).

#### Focused ultrasound and sonogenetic stimulation

The focused ultrasound (FUS) device consisted of two parts, the FUS transducer and an adapter. The FUS transducer was made of a lead zirconate titanate (PZT) ceramic resonator (DL-43, DeL Piezo Specialties) encapsulated by a 3D-printed housing (photopolymer resin, Formlabs). The PZT ceramic resonated at a frequency of 1.5 MHz and had an aperture of 15 mm and a radius of curvature of 25 (animal 1, animal 3) or 35 mm (animal 2). The FUS transducer was inserted into a 3D-printed adapter which allowed the FUS transducer to stably sit in the recording chamber. The FUS device was calibrated by a hydrophone (HGL-200, Onda).

Prior to FUS stimulation, the craniotomy chamber was thoroughly irrigated and rinsed with 0.9% saline, 10% povidone-iodine (Betadine), and Dakin’s solution (1:10 bleach solution in 0.9% saline). The recording chamber and the surface of the FUS transducer were filled with steam-sterilized, degassed ultrasound gel (Aquasonic 100, Parker). The FUS device was then inserted and secured to the craniotomy chamber. The FUS device was connected to a power amplifier (350L, Electronics and Innovation) driven by a function generator (33510B, Keysight). Similar to the electrical stimulation experiment, a TTL-pulse from the Datapixx would trigger the function generator to send a FUS stimulation (1.5 MHz, 10 HZ PRF, 50% duty cycle, 15s stimulation duration, acoustic pressure 1.0 MPa for animal 2, 1.2 MPa for animal 1 and animal 3; **Fig. S12a**) at a randomly selected inter-stimulation interval between 100 and 185 s. The ultrasound pressures correspond to mechanical indices of 0.82 and 0.98, respectively.

#### Pharmacological inactivation

Muscimol (GABA-A receptor agonist) was used to inactivate the hypothalamus during sonogenetic stimulation. 5 mg/mL muscimol (Sigma-Aldrich) in water was injected to the LHA. Muscimol (0.3–0.6 µL) was infused into LHA using a 32-or 64-channel linear array with a fluid capillary at a rate of 0.1-0.13 µL/min.

During LHA inactivation, the a12 ultrasound parameters were mostly the same as those used for the other sonogenetic experiments, except animal 1 ultrasound pressure was 1.4 MPa for pre-and post-muscimol injection comparisons. The baseline heart rates during sonogenetic stimulation pre-and post-muscimol infusion were used to compute the effect of LHA inactivation on heart rate.

#### Task design

Monkeys were trained to perform a visual decision-making task with offers of varying reward amounts, reward uncertainties, delay periods, and delay uncertainties (**Fig. 5a, b**). Each offer consisted of two vertical bars, one indicating the reward component and the other indicating the delay component. The reward component was displayed as a black vertical bar which could have short, medium, or tall height, respectively corresponding to juice reward amounts that were small, uncertain with P(large) = P(small) = 50%, or large. The reward component could also be displayed as a medium blue vertical bar indicating a 100% probability of receiving a medium reward amount. The delay component was displayed as a gray vertical bar which could have short, medium, or tall height, respectively corresponding to delay durations that were short, uncertain with P(long) = P(short) = 50%, or long. It could also be displayed as a medium yellow vertical bar for 100% medium delay duration. Thus, rewards and delays varied in their magnitude and uncertainty based on the heights and colors of their respective vertical bars. Finally, for each offer, the positions of the two components on the screen from left to right indicated the sequence in which they would be delivered. They could be either (1) reward first then delay, or (2) delay first then reward. The reward amounts corresponding to small and large settings were approximately 0.17 and 0.64 mL, while the medium reward amount was set to the average of the small and large amounts. The delay durations corresponding to small and large settings were 2 and 8 s, while the medium delay duration was set to the average of the small and large durations.

Two versions of the task were used: one used for neural recording and chemogenetic manipulation, and another for task performance manipulation with sonogenetics. For the neural recording and chemogenetic version of the task, the task began with a fixation dot located in the center of the screen, and the monkey needed to fixate for the trial to continue. Two offers were sequentially presented. The two offers were each randomly generated with the constraint that they could not be identical to each other. The monkey needed to choose from the two offers by gazing at the chosen offer for 0.5 s. Afterwards, the unchosen offer disappeared from the screen, and the task proceeded based on the offer the monkey chose. If the monkey chose an offer in which the delay was first, then the delay duration was revealed after 1 s (delay epoch). After an additional 1 s, the timer would begin lasting the duration of the delay. After the timer ended, the reward was revealed (reward epoch), with the reward delivered after another 1 s. If the monkey chose an offer in which the reward was first, then the reward was revealed after 1 s. After an additional 1 s, the reward was delivered. The delay was then revealed, and the timer would begin counting down the duration of the delay after another 1 s. The inter-trial interval (ITI) was set as 1 s.

For task performance manipulation via sonogenetics, the ITI was extended to 10 s to account for the 15 s ultrasound stimulation duration. The ultrasound parameters were almost the same as those used during sonogenetic stimulation without a task. However, for animal 2, the ultrasound pressure was lowered from 1.0 MPa to 0.87 MPa to prevent the monkey from aborting the trial. The ultrasound stimulation was applied starting halfway during the ITI, so the trial was initiated 5 s after ultrasound stimulation onset. Stimulation was randomly assigned to a trial with 30% probability. The task was also designed so that no two consecutive trials were designated as “stimulation trials” to limit the amount of ultrasound delivered to the brain in a short amount of time. For example, if one trial was designated a “stimulation trial,” then the next trial would have a 0% chance of being a “stimulation trial,” and the next trial after that would have a 30% chance of being a stimulation trial.

#### Neural recording

Single-unit recording was performed using 32-or 64-channel linear arrays (v-probes, Plexon). Signal acquisition (including amplification and filtering) was performed using a Plexon 40 kHz recording system. In these experiments, action potential waveforms were identified online by multiple time-amplitude windows, and the isolation and sorting were performed offline to identify clusters from the recording data (Kilosort2). Isolated clusters were further refined via a manual curation process to identify clusters corresponding to well-isolated single neurons.

#### Neuroimaging

Anatomical images of the monkey brain were acquired via magnetic resonance (MR) imaging. At the time of MR imaging, monkeys were anesthetized, transported to the MR scanning room, and placed in the supine position. The monkey’s head was placed in a knee coil (Siemens) and inserted into the MR scanner (3T, Siemens). Whole brain images were acquired using the T1-MPRAGE sequence averaged three times by the scanner. Occasionally, the monkey brain was scanned with a microelectrode in the brain to verify the stimulation and/or recording location (**Fig. S2a**). For this procedure, microelectrodes were inserted into the monkey brain and lowered to the target region (details in *Electrical stimulation* and *Neural recording*). The monkey brain was then imaged in the MR scanner using the previously described sequence.

MR thermometry was used to noninvasively quantify the temperature rise associated with FUS stimulation in area 12. At the time of MR imaging, monkeys were anesthetized and head fixed to an MR-compatible stereotaxic frame via ear bars and a bite bar in the prone position (Kopf Instruments). Similar to the FUS stimulation in awake monkey procedure, the recording chamber was filled with degassed, sterilized ultrasound gel, and the FUS device was positioned and fixed to the recording chamber to target area 12. A loop coil (11 cm diameter, Siemens) was placed on top of the head encapsulating the recording chambers to facilitate a12 imaging. The entire setup was inserted into the MR scanner. Temperature maps were generated with a continuously applied gradient-echo imaging sequence with a flip angle of 20 degrees, TR of 107 ms, TE of 12 ms, slice thickness of 3 mm, and a matrix size of 384×384 for a 173×173 mm field of view. Magnitude and phase images were processed using ThermoGuide software (Image Guided Therapy) to generate temperature maps. FUS was applied during MR scanning to determine the optimal parameters for increasing the local brain temperature to drive TRPV1 activation in a12. After each stimulation, T2-FLAIR scans were acquired to assess any tissue damage.

#### Histology

After all experiments were complete, animals were deeply anesthetized with sodium pentobarbital and transcardially perfused with 1X PBS, followed by 4% paraformaldehyde (PFA) in 1X PBS (pH 7.4). The brain was postfixed in 4% PFA overnight, blocked, and equilibrated with 30% sucrose in 1X PBS at 4°C. Frozen sections were then cut on a sliding microtome at 50 µm thickness in the coronal plane and stored in 1X PBS with 0.1% sodium azide at 4°C in a 12-well plate. To visualize the immunoreactive signals of green fluorescent protein (GFP) co-expressed with hM4Di, we followed the methodology described previously (*47*). Sections of ROIs were immersed in 1% skim milk containing 0.1% TritonX-100 for 1 hour at room temperature and washed 1x with a blocking buffer (1x PBS containing 0.1% TX-100 and 1% normal goat serum) for 15 minutes at room temperature. The sections were then incubated overnight at 4 °C with rabbit anti-GFP monoclonal antibody (1:500, Thermo Fisher Scientific, G10362) in blocking buffer for 2 days at 4°C. The sections were washed 3×10 minutes with 1x PBS and then incubated in the same fresh medium containing biotinylated goat anti-rabbit immunoglobulin G antibody (1:1000, Jackson ImmunoResearch) for 2 hours at room temperature, washed 3×10 minutes with 1x PBS, incubated in avidin-biotin-peroxidase complex (ABC Elite, Vector Laboratories) for 2 hours at room temperature, and washed another 3×10 minutes with 1x PBS. To visualize GFP, the sections were reacted in 3,3-diaminobenzidine (DAB) with cobalt chloride and nickel chloride metal enhancers (ThermoFisher, #34065). The sections were mounted on gelatin-coated glass slides, air-dried, dehydrated in an ethanol series, cleared with xylene, and cover-slipped with DPX (#06522, Sigma). Sections were imaged with standard brightfield microscopes (Keyence, Nikon AXR) to visualize the signals.

Fluorescent in situ hybridization (FISH) staining using the RNAscope assay was performed following the manufacturer’s protocol for RNAscope Multiplex Fluorescent v2 kit adapted for large, 50 µm thick slices (ACD). The RNAscope probes used for visualizing TRPV1, were Rn-Trpv1-C3 (ACD, #501161-C3), Mm-Slc17a6-C2 (ACD, #319171-C2), and co-stained with DAPI mounting medium (Vector Labs). Sections were imaged with standard fluorescent microscopes (Keyence, Zeiss LSM 980 Airyscan 2) to visualize the signals.

### Data analysis and statistical analysis

#### Heart rate metrics quantification

PPG signals were aligned to stimulation onset via TTL pulse timing and high-pass filtered (5th-order Butterworth, 1 Hz cutoff, zero-phase forward-backward filtering) to remove low-frequency drift. Beat onsets and peaks were detected using the PPG-beats MATLAB toolbox (*102*); saturated segments were corrected via cubic spline interpolation (*103*). Heart rate within a time window was computed as beat count divided by window duration, with fractional interbeat intervals included at the window edges. Heart rate was expressed as percent change from baseline (mean heart rate, 5 s pre-stimulation). For time course visualization, the instantaneous heart rate (inverse of the interbeat interval) was assigned to the later beat of each pair and linearly interpolated to the PPG sampling rate.

Stimulation sites were screened for heart rate modulation by computing HR change 0-1.5 s post-stimulation onset and testing for a significant difference from zero (signed-rank test, p < 0.05). The electrical stimulation did not evoke movement (**Fig. S4**). However, occasionally, the animals happen to be moving by chance. To make sure that such occasional movement did not pollute our PPG results, we excluded timeseries using a movement index (number of peaks in high-pass-filtered force-transducer signal captured 5 s before and after stimulation). The data were excluded if it contained >15 peaks.

The trials at each site were used to classify the site as producing a significant HR increase, decrease, or no effect, forming the basis for pooled “effective site” and “ineffective site” responses (**Fig. 1c**, **1k**). For cross-validated analyses (**Fig. 1d**, **1m**), quantification and visualization were used with half of the stimulations, with statistical analysis computed from the other half of stimulations.

Heart rate variability (HRV) was defined as the standard deviation of interbeat intervals within a given window and expressed as percent change from baseline (5 s pre-stimulation). The HRV time course was computed via 3 s moving standard deviation of the interbeat-interval time course, with each value assigned to the window’s end time. Mean HRV changes were calculated by averaging the percent-change time course over early (0-1.5 s) and late (4-5.5 s) post-stimulation windows.

For chemogenetic inactivation experiments, PPG signals were processed as above to obtain heartbeat peaks on a session-by-session basis. Each session was divided into pre-and post-injection periods and split into 30 s windows; windows with high movement indices (>10 peaks) were excluded. Each post-injection window was treated as a separate measurement, with heart rate change computed relative to the mean pre-injection heart rate for that session. Rank-sum tests compared heart rate changes between DCZ and saline injections in a12 and LHA.

For sonogenetic stimulation experiments, PPG signals were processed similarly on a stimulation-by-stimulation basis and aligned to stimulation onset via TTL timing. Stimulations were excluded for high movement index (>15 peaks, −15 to 60 s relative to onset), and analyses were restricted to low-arousal trials. Specifically, for animal 1, k-means clustering (k=2) of pre-stimulation baseline heart rate identified “low” and “high” states, and only “low” trials were retained; all trials were used for animals 2 and 3. Heart rate and HRV time courses were computed using 10 s windows: heart rate as the inverse of the mean interbeat interval per window, expressed as percent change from the 30 s pre-stimulation baseline; HRV as the standard deviation of interbeat intervals per window, expressed as percent change from the pre-stimulation baseline HRV. Sonogenetic stimulations during muscimol inactivation were processed in the same way.

To evaluate the impact of muscimol inactivation of LHA on heart rate and HRV, pre-and post-injection heart rate traces were acquired using 30 s windows before sonogenetic stimulation, with heart rate changes computed as in the chemogenetic experiments and compared directly to saline infusions in LHA from the chemogenetic dataset. To evaluate the impact of muscimol inactivation of LHA on sonogenetic-evoked responses, HRV values pre-and post-injection were averaged 0-30 s post stimulation window (three 10 s time bins), the period capturing most of the sonogenetic-evoked responses prior to muscimol infusion.

#### Electrical stimulation threshold, latency, and arousal state analyses

We compared the heart rate decrease effects of a12 and LHA stimulation in terms of current sensitivity, latency, and dependency on arousal state (**Fig. 3a-c**). For current sensitivity, sites were included if they elicited a significant heart rate decrease at −300, −200, −100, or −50 μA using the same heart rate computation method. Stimulations at currents that did not evoke a significant effect (i.e. a12 stimulation at −50 μA) were pooled by their −300 μA site classification. For latency, “effective” stimulations from a12 and LHA were compared. Latency was defined as the time post-stimulation at which the z-scored signal (computed with 1.5 s pre-stimulation baseline) first reached ≥4 and sustained that level for ≥100 ms. For arousal state, each animal’s pre-stimulation baseline heart rate was used as a proxy for arousal state (*42*), with “low” and “high” states defined as the bottom and top 25% baseline heart rates, respectively.

#### Auxiliary behavior quantification

Movement signals (see above) were converted to percent change time courses using a 5 s pre-stimulation baseline. The percent change within a window was the mean signal over that window (**Fig. S4**). For eye-closed state analysis, the analog pupil signal was processed similarly, then binarized (eyes closed = 0, open = 1) using a pupil threshold. Blink artifacts (trains of 0’s <500 ms) were removed. “Eye-closed” stimulations were defined as having eyes closed from −1 to 1.5 relative to stimulation onset, and their heart rate change effects were analyzed separately (**Fig. S5**).

#### Neuronal data

We recorded a total of n=293 neurons in area 12 (animal 1, n=219, animal 2, n=74). We analyzed the subset of neurons that were recorded during both task performance and during resting state, which were recorded for a total duration of at least 500 seconds during resting state during which simultaneous heartbeat recordings were available. This resulted in a database of n=108 neurons (animal 1, n=88, animal 2, n=20) that were recorded during resting state for an average of 988 s (standard deviation = 375 s, range = 685-1877 s).

#### Power spectral analysis of resting state spiking activity and heartbeats

We first computed the cross-correlation between each neuron’s spike train and its analogous heartbeat train. These signals were initially represented as binary vectors (1 for a spike or heartbeat, 0 otherwise) with a temporal resolution of 1 ms, then were multiplied by 1000 to convert from units of spikes/ms (or beats/ms) to conventional units of spikes/s (or beats/s), and the cross-correlogram spanned the range of temporal lags from −1500 ms to +1500 ms. To control for very slow drifts in spiking and/or heart rates, we used linear regression to regress out simple slow effects of time from each signal (using regressors representing a constant factor and the linear, quadratic, and cubic effects of time), thus obtaining a residual spiking signal and residual heartbeat signal. Then we computed the cross-correlogram between the residual spiking and residual heartbeat signals. To analyze their relationship in the frequency domain, we computed the cross-correlogram’s power spectral density (sampling with resolution of 0.33 Hz), and as a summary measure, computed the total power spectral density in each of three frequency bands (0.3-2 Hz, 2-4 Hz, and 4-6 Hz, representing bands below, overlapping, or above the observed heart rates in this data).

We next tested whether each neuron’s power spectral density was different from what we would expect by chance, under the null hypothesis that the spiking and heartbeat signals both had the temporal structure that we observed in the real data but had no true temporal coordination with each other. To do this, we used non-parametric tests based on randomly time-shifting the data. For each neuron, the above analysis was repeated on n=100 circularly shifted datasets, in which the heartbeat data vector was circularly shifted by a random temporal offset, which was drawn uniformly from the set of all possible time shifts with 1 ms resolution (subject to the constraint that the absolute value of the offset was > 20 s, to ensure the offset was large enough to meaningfully shift the temporal relationship between spiking and heartbeats). We then subjected each shifted dataset to the same analysis as above. We defined a neuron as significantly temporally related to the heartbeat in a given frequency band at *α* = 5% (two-tailed test), if the power spectral density in that band from the real data was more extreme (either higher or lower) than 97.5% of the analogous measures from the shifted data.

To test whether neuronal activity changes occurred more prominently before or after heartbeats, we performed a second analysis of the cross-correlograms between residual spiking and residual heartbeats, computing a “spike-heart order index”. We defined the pre-heartbeat activity, *x_before_*, as the mean of the first half of the cross-correlogram (lags from −1500 ms to −1 ms), and post-heartbeat activity, *x_after_*, as the mean of the second half (lags from +1 ms to +1500 ms). We then defined the order index as:

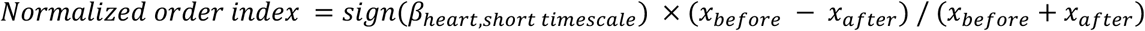

Where the sign(*β_heart,short timescale_*) term normalizes the index so it aligns with the direction of the neuron’s heart-related activity, based on the sign of the fitted effect of short timescale changes in heart rate on the cell’s activity (from the model described below). That is, if a cell changes its activity more before than after heartbeats, then it has *x_before_* > *x_after_* if its activity is positively related to heart rate (sign = +1), but has *x_before_* < *x_after_* if its activity is negatively related to heart rate (sign = −1). In addition, we aimed to ensure that each neuron’s order index was scaled based on the magnitude of that neuron’s rate modulation relative to its mean firing rate (e.g. a neuron with 1 Hz pre-vs-post modulation should have a larger index if its mean firing rate is 1 Hz than if it was 10 Hz). This would not occur if we used the residual cross-correlograms directly, because the regression included a constant factor representing the mean firing rate, causing the residuals to have zero mean. Therefore, before doing the above analysis, for each cell, we applied a vertical offset to the residual cross-correlogram to set its mean equal to the mean of the original (raw) cross-correlogram. To test for significance, we repeated the above analysis on n=100 circularly shifted datasets. We defined a neuron as having a significant normalized spike-heart order index at *α* = 5% (two-tailed test), if the index from the real data was more extreme (either higher or lower) than 97.5% of the analogous indices from the shifted data.

#### Model of resting state heartbeat-related activity

To model the relationship between spiking activity and heartbeats, we fit each neuron’s spike probability at each millisecond during resting state using a simple linear regression model, as a weighted linear function of eight variables: (1) a constant factor, (2) local heart rate estimated on the timescale of seconds, obtained by smoothing the heartbeat vector with a Gaussian kernel with σ = 2 s, (3) local heart rate estimated on the timescale of minutes, by smoothing with σ = 60 s, (4-5) same as 2-3, but for eye open/closed state, based on an analogous binary vector representing whether the eyes are open at each millisecond (1 for eyes open, 0 otherwise), (6-8) linear, quadratic, and cubic effects of elapsed time during the session.

To test whether a model indicated a significant temporal coordination between spiking activity and heartbeats, we repeated the above analysis on n=100 circularly shifted datasets in which a random temporal offset was used to shift the heartbeat train, eye state, or both (analogous to the procedure described above for power spectral analysis). We defined the model fit to a neuron as significantly temporally related to the heartbeat at *α*= 5% if the deviance of the fit to the real data was lower than the deviance of 95% of the analogous measures from the heartbeat-shifted data (one-tailed test, because randomly shifting the data can in expectation worsen the fit but not improve it). We defined a fitted regression weight as significant if it was more extreme (higher or lower) than 97.5% of the analogous fitted regression weight from the heartbeat-and-eye-shifted data (two-tailed test). We also found a highly consistent pattern of results using an alternate statistical approach, based on comparing the deviance or regression weight from fits to the real data against surrogate datasets where the heart, eye, or both from the session were replaced by the corresponding heart, eye, or both signals from each of the other experimental sessions (**Fig. S7**).

#### Analysis of neural activity during the task

We analyzed each neuron’s firing rate in a pre-choice time window (0.8 s before choice). We quantified the neuron’s relationship to motivational task variables by using a linear regression model to fit firing rate as a weighted linear function of the following regressors based on the features of the chosen offer: (1) a constant factor, (2) offer type (1 if reward-first, 0 if delay-first), (3-8) the following offer attributes only for reward-first offers: expected reward magnitude, expected time delay, reward uncertainty (1 if uncertain, 0 otherwise), delay uncertainty, interaction between expected reward magnitude and delay, interaction between reward uncertainty and delay uncertainty, (9-14): those same attributes but only for delay-first offers, (15) a binary regressor indicating which offer was chosen (0 for offer 1, 1 for offer 2). A neuron was identified as having significant task-related activity using a standard one-tailed chi-squared test of the deviance (i.e. whether the deviance of the model fit including all terms was significantly lower than the deviance of a null model fit that only included the constant factor). P-values for individual fitted regression weights were determined using standard two-tailed t-tests.

We then sought to quantify how heart-related and motivation-related activity were related to each other. In particular, are they unified reflecting a single unified signal transmitted by the same subpopulation of neurons? Or are they separable, such as being transmitted by uncorrelated or even entirely distinct subpopulations of neurons? To do this, we used a modeling approach to quantify their relationship while controlling for the statistical false positive and false negative rates in the dataset. This is important because even if the “unified” hypothesis was completely correct, we would still expect some individual neurons to have only one signal significant (or to have no signals significant) due to statistical false negatives; similarly, even if the “distinct” hypothesis was completely correct, we would still expect some individual neurons where both heart and motivation effects reach significance due to statistical false positives.

We therefore used a two-stage modeling approach. In the first stage, separately for each signal type *s* (heart or motivation), we fit a model of the probability of detecting a significant signal as a cumulative gaussian function of the number of samples of data *n* for a given neuron:

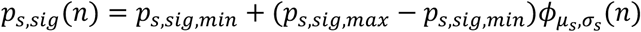

This model had three parameters: μ*_s_*, the mean of the gaussian; σ*_s_*, the SD of the gaussian, and *p_s,sig,max_*, the maximum probability of significance (occurring at the maximum sample size in the dataset). The fourth term *p_s,sig,min_* is the minimum probability of significance and was fixed at 0.05, because that was the nominal false positive rate of the statistical tests for each signal. These parameters were all constrained to be >= 0 and the maximum likelihood fits were obtained using the constrained nonlinear multivariable function solver in Matlab (‘fmincon’). Given these parameters for a given signal, we can obtain an estimate of the fraction of neurons that truly transmit that signal *p_s,true_*, by considering that for neurons with the maximum number of data points, their corresponding maximum probability of significance is the result of a mixture of neurons with true signals and neurons with false positives (occurring at a rate equal to *p_s,sig,min_*).

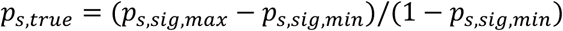

This produced estimates of *p_heart,true_* = 0.45 and *p_motiv,true_* = 0.15. In the second stage, having modeled each signal individually, we then modeled how heart and motivational signals co-occurred across the population. We characterized the coupling between these signals using a single key parameter *p_both,true_*: the probability of a cell having true signals related to both heart and motivation. We computed the most extreme values this parameter could take, corresponding to the “distinct” and “shared” hypotheses, as well as the intermediate value it could take corresponding to the “uncorrelated” hypothesis, all subject to the constraint of matching the estimated marginal probabilities from the first stage model (*p_heart,true_* and *p_motiv,true_*):

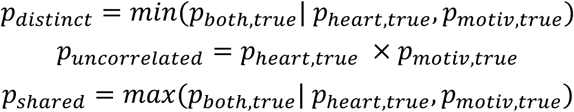

We then used these as reference points to define the assortment index, so the index was −1, 0, or +1 when *p_both,true_* was equal to *p_shared_*, *p_uncorrelated_*, or *p_distinct_*, respectively, and was linearly interpolated when *p_both,true_* was in between those reference points:

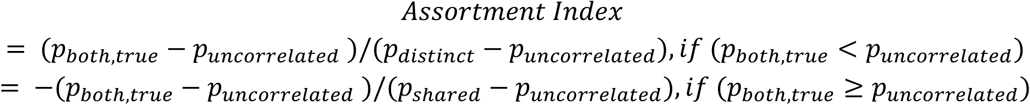

We then estimated the assortment index using a maximum likelihood fit to the a12 population. This fit was done using the constrained nonlinear single-variable function solver in Matlab (‘fminbnd’), constraining the index to be in the bounds [-1, +1], and enforcing the constraints of the first stage models mapping each neuron’s sample size onto a detection probability for each signal. This produced a best fitting assortment index of 0.40. To validate our approach, we generated n=400 simulated datasets for each assortment index in the set {-1, −0.8, −0.6, −0.4, −0.2, 0, +0.2, +0.4, +0.6, +0.8, +1}. Each simulation was constrained to generate each neuron’s significance for each signal based on the simulation’s specified assortment index (and its corresponding *p_both,true_*), but was otherwise as closely matched as possible to the real dataset (i.e. using the first stage models to set *p_heart,true_*, *p_motiv,true_*, and each individual neuron’s estimated detection probability for each signal based on that neuron’s sample size). We defined the real data as significantly different from a given simulation at *α*= 5% if the fitted assortment index to the real data was more extreme (either higher or lower) than 97.5% of the distribution of the fitted assortment indexes to the n=400 simulated datasets.

To further validate this conclusion, we examined the relationship between heart-and motivation-related activity using additional approaches, which all produced consistent results (**Fig. S14**).

First, we repeated our GLM analysis of each neuron’s firing rate in six different time windows during the task: offer 1 (0.1-0.5 s after offer 1 onset), offer 2 (0.1-0.5 s after offer 2 onset), pre-choice (0.8 s before choice), pre-reward reveal (1 s before outcome reveal), pre-delay reveal (1 s before delay reveal), and pre-timer end (1 s before timer end). In the two offer windows, we used the same linear regression model as above but only as a function of the features of the presented offer, and we did not include the regressor for which offer was chosen since the second offer had not been presented yet. In the other windows, we used the same regressors but as a function of the features of the chosen offer. In each window, we tested whether significant task-and heart-related activity co-occurred in single cells more often than chance, against the null hypotheses that these two properties were statistically independent of each other. To do this, we tested whether the fraction of cells where both were significant (*p_both_*) was different from the fraction expected by chance if the two were independent (*p_heart_* x *p_motiv_*; two-tailed binomial test). In all cases the fraction was very close to chance level with no significant difference (**Fig. S14**), with the fraction tending to be slightly below chance, consistent with separable signals represented in a manner between the “uncorrelated” and “distinct” hypotheses. Second, we focused on the specific set of motivational variables in each time window that were encoded by the population, and asked whether such coding was associated with significant heartbeat-related activity. To test this, we repeated the above test for each individual regressor weight from the model of task-related activity, replacing *p_motiv_* with *p_motiv,regressor_*, i.e. the fraction of the population with a significant weight for that individual regressor. Again, the co-occurrence of heart-and motivation-related signals was close to chance levels for every motivational variable in every task epoch, with none being significantly different from chance (**Fig. S14**).

## Acknowledgments

This work was supported by the National Institute of Mental Health under award numbers R01MH128344, R01MH110594, and R01MH116937; the Conte Center on the Neurocircuitry of OCD MH10643 to I.E.M; and Brain Initiative Grant to HC (with IEM) UG3MH126861. We thank Ms. Kim Kocher for excellent animal care and Dr. Joel Price for sharing his anatomy collection. We also thank Dr. Suzanne N. Haber for sharing her anatomy collection and allowing us to examine related anatomical cases. We are grateful to Ms. Rebecca Mellor and Dr. Andreas Burkhalter from the Washington University Histology Innovation and Service Center (WHISC) for their assistance in sectioning and processing brain tissue. We also thank Dr. Peter Bayguinov from the Washington University Center for Cellular Imaging (WUCCI) for imaging brain sections. We acknowledge Dr. Lawrence Snyder for insightful discussions during the early planning of sonogenetic experiments in non-human primates. We also acknowledge the Hope Center Viral Vector Core and the Hope Center for Neurological Disorders for synthesizing viral vectors for some of these studies. Lastly, we acknowledge members of the Monosov and Chen labs for many insightful discussions.

## Author contributions

KX and IEM designed the research. IEM guided the research. SPE and IEM designed the decision-making task. KX, TO, MY, and AHS implemented the experiments and collected data. JY conceived a novel in situ hybridization protocol for monkey brain tissue processing. KX analyzed the heart rate data. ESBM analyzed the neuronal activity and decision-making data sets with input from IEM. TM and KI assisted with implementation of chemogenetic viral vectors. SNH provided additional tracer injection case studies and advice. KX and IEM prepared the manuscript, and ESBM provided valuable comments. HC advised on ultrasound design and utilization, provided ultrasound equipment, and made the virus used in sonogenetics experiments. HC and IEM procured funding for this study. All authors contributed to the final version of the manuscript.

## Data availability

The processed data supporting the findings of the study will be made available upon publication. All original code will be deposited to GitHub upon publication. Any additional information is available from the lead contacts upon reasonable request.

