## Supplemental Material and Figures for "A prefrontal cortex-hypothalamus circuit for heart rate control in non-human primates"

### Figure S1

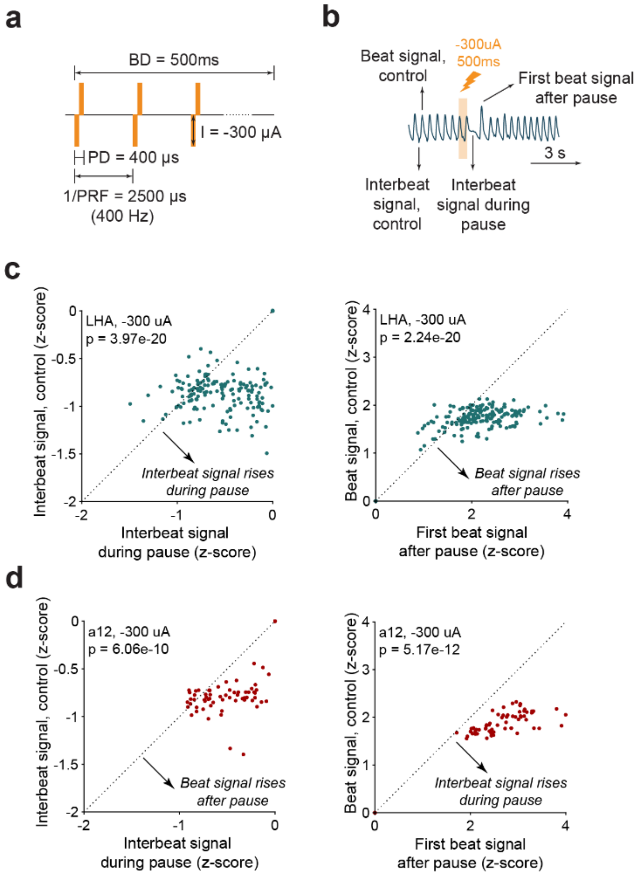


**Figure S1: Further characterization of photoplethysmography (PPG). a,** Schematic illustration of the electrical stimulation waveform. **b,** Schematic illustration of terminology used for PPG analyses, defining the “beat signals” and “interbeat signals” during “control” periods and periods right after stimulation. These metrics were quantified and compared to demonstrate that the PPG signal during the pause is not a result of signal drop-out. **c,** PPG beat analysis of electrical stimulation in effective sites within LHA. Left: Scatter plot comparing the beat signals between stim and control periods. Right: Scatter plot comparing interbeat signals between stim and control periods. **d,** PPG beat analysis of electrical stimulation in effective sites within a12. Left and right plots follow the same format as (c).

### Figure S2

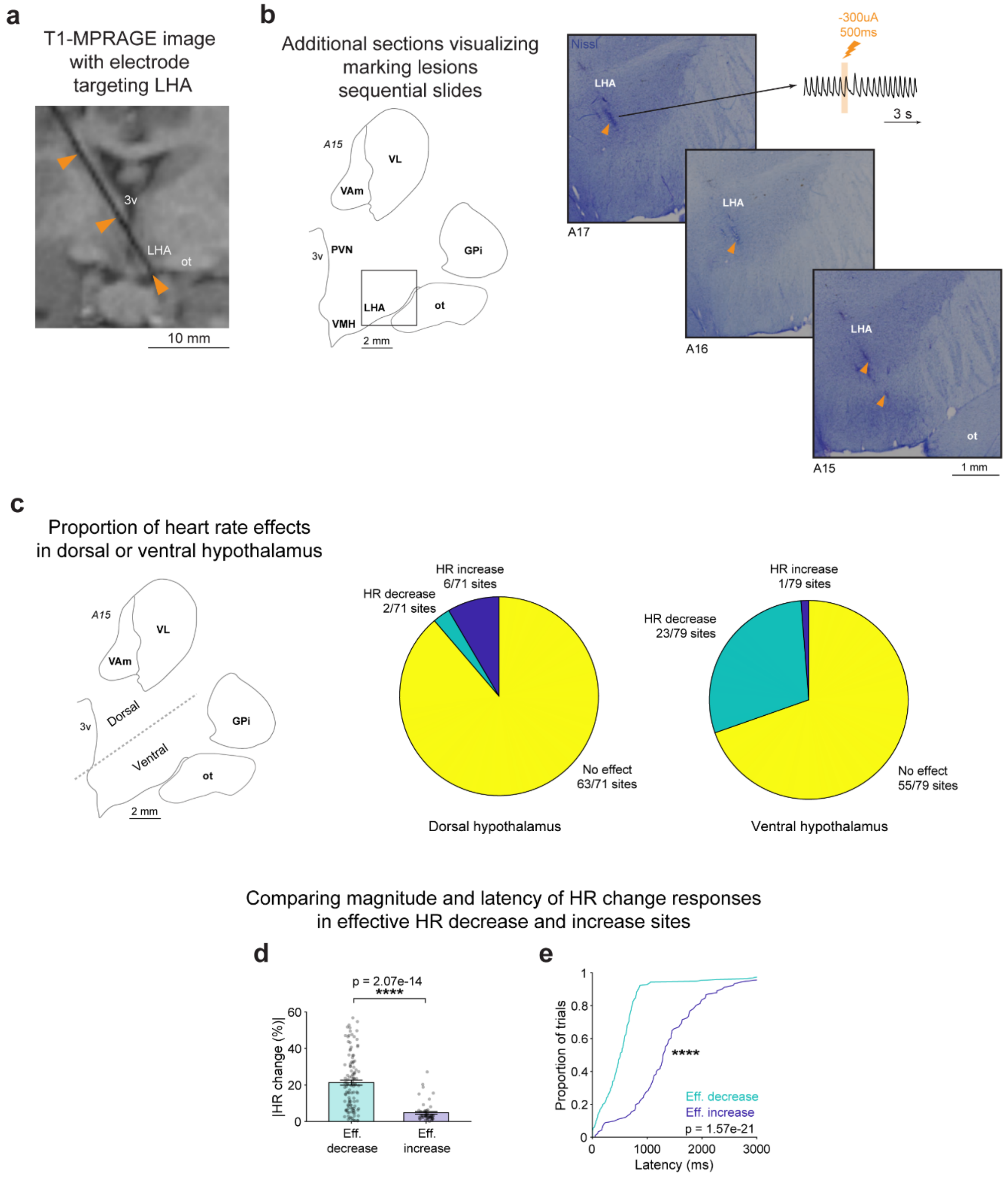

a
T1-MPRAGE image with electrode targeting LHA
3v
LHA
ot
10 mm
b
Additional sections visualizing marking lesions sequential slides
A15
VL
VAm
PVN
3v
GPi
LHA
ot
VMH
2 mm
Nissl
LHA
-300uA
500ms
3 s
A17
A16
A15
ot
1 mm
c
Proportion of heart rate effects in dorsal or ventral hypothalamus
Dorsal
Ventral
HR increase 6/71 sites
HR decrease 2/71 sites
No effect 63/71 sites
Dorsal hypothalamus
HR increase 1/79 sites
HR decrease 23/79 sites
No effect 55/79 sites
Ventral hypothalamus
Comparing magnitude and latency of HR change responses in effective HR decrease and increase sites
d
p = 2.07e-14
****
|HR change (%)|
Eff. decrease
Eff. increase
e
Proportion of trials
Latency (ms)
Eff. decrease
Eff. increase
p = 1.57e-21

**Figure S2: Imaging visualization of LHA targeting. a,** T1-weighted MPRAGE image with electrode targeting LHA in the coronal and sagittal planes. **b,** Additional Nissl images showing marking lesions within LHA. Left: Schematic illustration of monkey brain in the coronal plane at interaural A15, with a box highlighting the estimated imaging region for locating the marking lesions. Right: sequential slides showing marking lesions in LHA. **c,** Proportion of sites within the dorsal and central hypothalamus evoking heart rate decrease and increase responses. Left, Schematic illustration of the monkey brain in the coronal plane at interaural A15. The dotted line corresponds to the self-defined dorsal-ventral threshold. Right, Pie charts showing the proportion of sites in dorsal and ventral sites that evoked significant heart rate increase effects, significant heart rate decrease effects, or no effects. **d,** Magnitude of heart rate change in effective hypothalamus sites with HR decrease and increase effects, quantified as the absolute value of the heart rate change. n = 116, 54 stimulations for effective HR decrease and increase sites, respectively; P = 2.07e-14 (two-tailed rank-sum test). **e,** Cumulative distribution function plot of electrical stimulation-evoked heart rate response latencies in effective hypothalamus sites with HR decrease (green) and HR increase effects (blue). **** indicates P < 0.0001.

### Figure S3

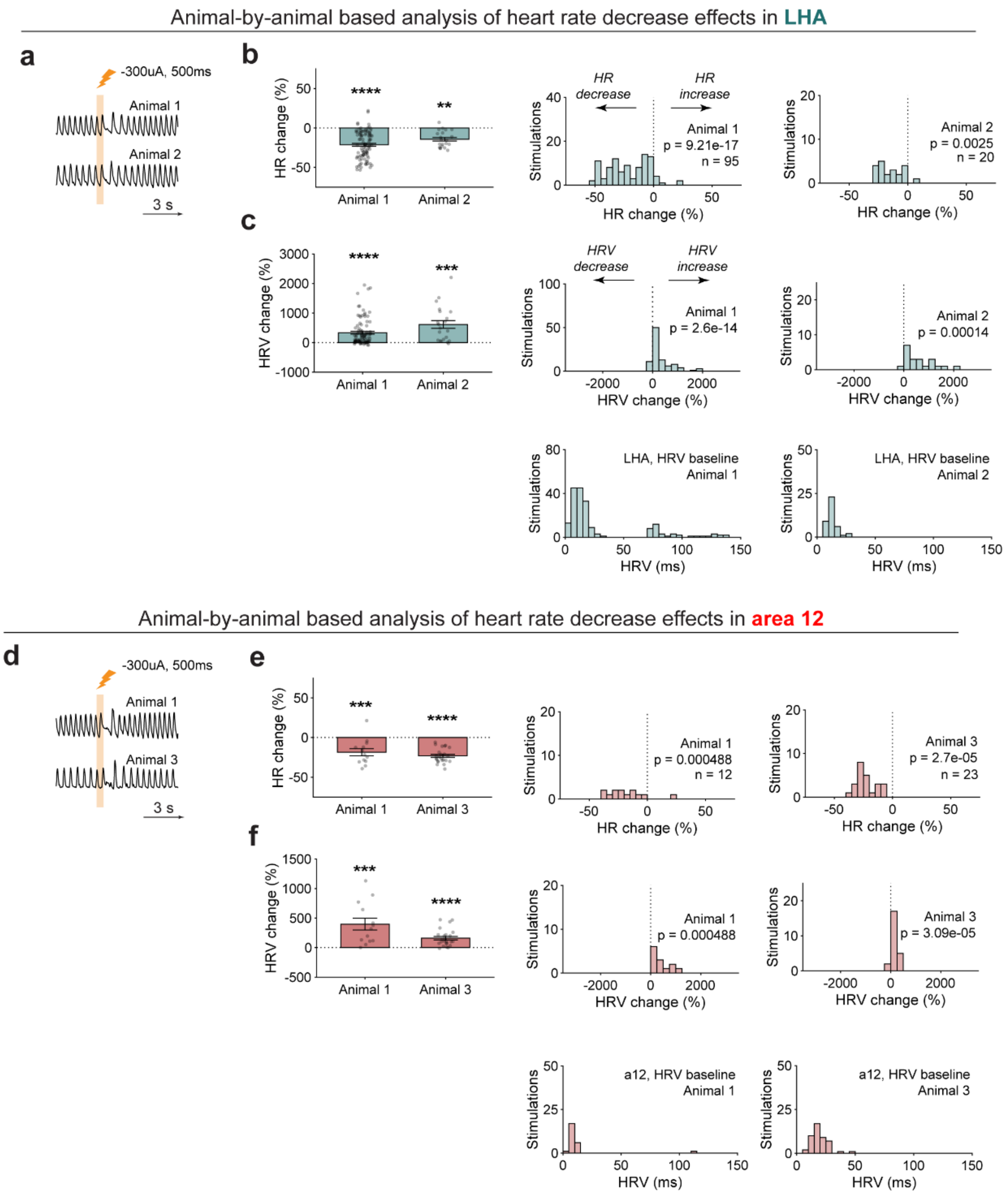


**Figure S3: Subject-based analyses of effective site electrical stimulation-evoked responses in LHA and a12. a,** Representative PPG traces for effective site stimulations in LHA across two animals. Yellow bar indicates electrical stimulation period applied at -300 μA and 500 ms. **b,** Heart rate responses for effective site stimulations in LHA across two animals. Left: Average heart rate change plotted for both animals. Data are mean ± s.e.m. Asterisks indicate signed-rank test against a hypothetical median of 0. Middle: Histogram of the heart rate change effects for animal 1. Right: Histogram of the heart rate change effects for animal 2. **c,** Heart rate variability responses for effective site stimulations in LHA across two animals. Left: Average heart rate variability change plotted for both animals. Data are mean ± s.e.m. Asterisks indicate signed-rank test against a hypothetical median of 0. Middle: Histogram of the heart rate variability change effects for animal 1. Right: Histogram of the heart rate variability change effects for animal 2. Histograms below show the baseline heart rate variability computed -5-0s relative to stimulation onset for each accompanying animal. **d,** Representative PPG traces for effective site stimulations in a12 across two animals. **e,** Heart rate responses for effective site stimulations in a12 across two animals. Left, middle, and right plots follow the same format as (b). Asterisks indicate signed-rank test against a hypothetical median of 0. **f,** Heart rate variability responses for effective site stimulations in a12 across two animals. Left, middle, and right plots follow the same format as (c). Asterisks indicate signed-rank test against a hypothetical median of 0. Histograms below show the baseline heart rate variability computed -5-0s relative to stimulation onset for each accompanying animal. Throughout this figure, *, **, ***, **** indicate $P < 0.05$, 0.01, 0.001, 0.0001, respectively; n.s., non-significant.

### Figure S4

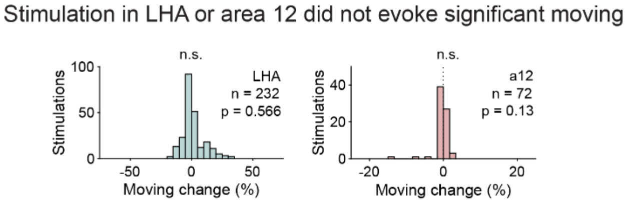


**Figure S4: HR-related site stimulation in LHA and a12 did not evoke movement.** Histogram of moving change evoked by effective site stimulation 0-1.5s after stimulation onset. Left: Histogram of effective site stimulations in LHA. Right: Histogram of effective site stimulations in a12. n.s. denotes non-significant (signed-rank test against a hypothetical median of 0).

### Figure S5

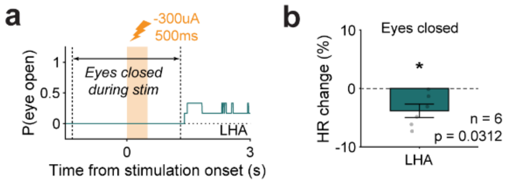


**Figure S5: Heart rate reduction effects during LHA stimulation when animal eyes are closed. a,** Probability of eye open time course to select stimulations with eyes closed during the stimulation period. **b,** Average heart rate change during eyes closed stimulations. Data are mean ± s.e.m. * above the bar indicates $p < 0.05$ (signed-rank test against a hypothetical median of 0).

### Figure S6

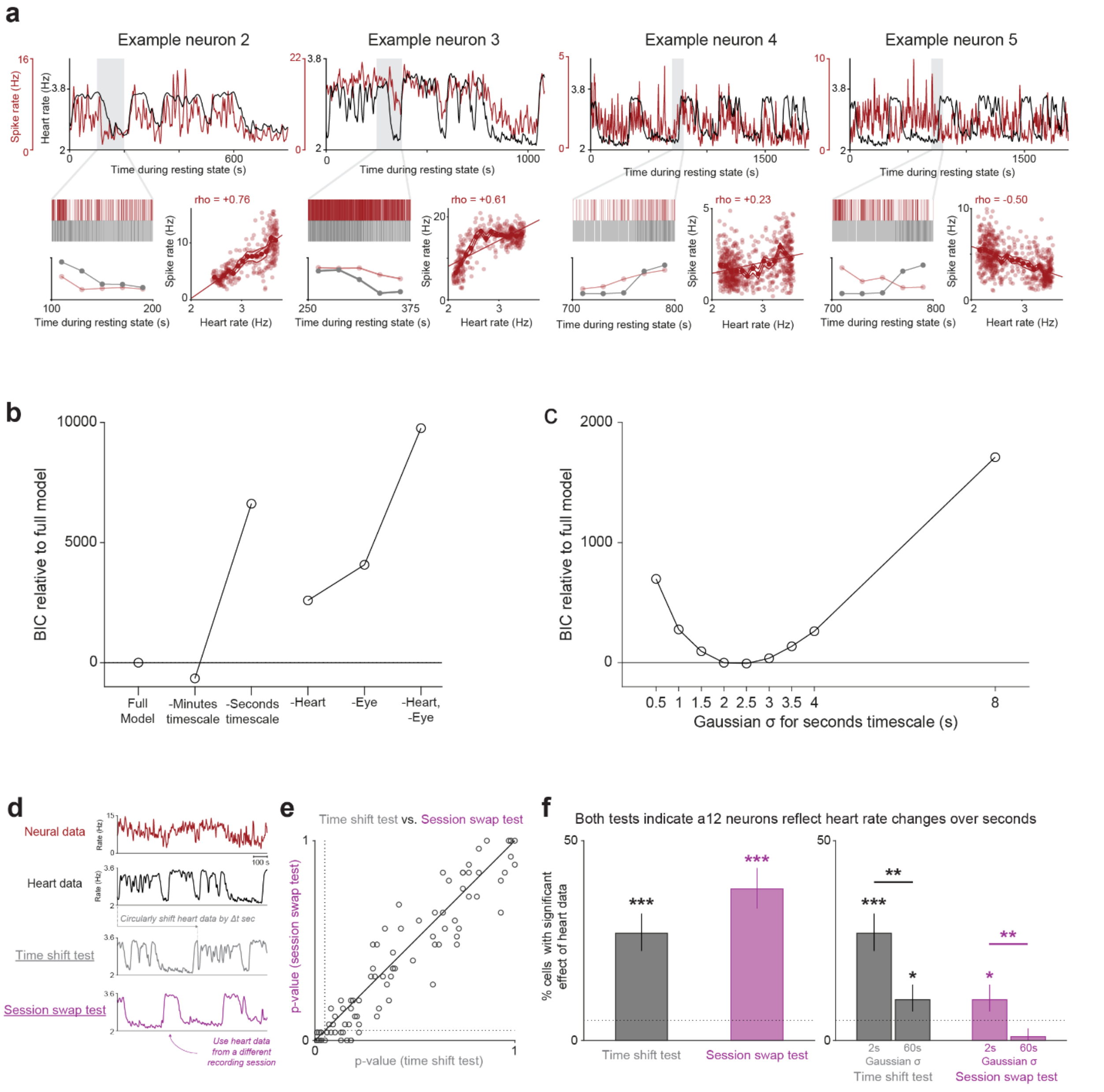


**Figure S6: Supplemental analyses demonstrating heart-related activity modulations on the timescale of seconds. a,** Additional example neurons illustrating relationships between spike rate and heart rate. Same format as main text. Neuron *2*: positive relationship for a neuron recorded in a different session from the main text example. Neuron *3*: positive relationship for a neuron recorded in the same session as the main text example. Neurons *4* and *5*: positive vs. negative relationships for two neurons recorded in the same session as each other. **b,** model comparisons demonstrating importance of seconds timescale for modeling neuronal activity. Shown are relative Bayesian information criterion (BIC (*96*); lower is better) of the full model used in the main text, and of alternate versions that remove minutes timescale regressors, seconds timescale regressors, heart regressors, eye regressors, or both heart and eye regressors. This suggests activity is best fit by modulations over the timescale of seconds, not minutes, indicated by the fact that removing the minutes timescale actually improves the BIC (i.e. lower than full model), while removing the seconds timescale greatly worsens the BIC. This also suggests activity is best fit by relationships with both heart and eye, indicated by the worsening of BIC if either is removed from the model. **c,** model comparisons demonstrating the best fitting seconds timescale appears to be approximately 2 seconds (same format as main text figure but expressed as BIC and with an expanded parameter space). **d-f,** Similar results using two distinct methods for statistical tests. **d,** The two methods. Shown are the example neurons from the main text's neural data (red) and heart rate (black). The main text reports results of a *circular shift test*: using n=100 random circular time-shifts of the heart rate (one example time-shift is shown in gray) to represent the null hypothesis that spiking and heartbeats are temporally unrelated to each other. Here we also show a *session swap test*: swapping out the real heartbeat vector from the neuron's session with the heartbeat vectors from n=25 other sessions (one example other-session heartbeat vector is shown in purple), representing the null hypothesis that spiking is no more related to heartbeats in its own session than other random sessions. **e,** both methods produced very similar results when testing the overall significance of neuron-heart relationships, indicated by great similarity between p-values from the time-shift test (x-axis) and session-swap test (y-axis). **f,** both methods produced qualitatively similar results. The session-swap test was somewhat more sensitive to overall effect of heart data (left) and somewhat less sensitive to effects of each specific heart regressor (right), likely due to the discretization of p-values (due to the limited number of other sessions being available for the swap test). However, they both produced the same overall pattern of results: a large fraction of neurons having significant relationships between spiking and heartbeats (left), an above chance number of them having significant relationships specifically with seconds timescale heart rate (right, 2 s bars), and significantly more neurons with significant effects of seconds timescale than minutes timescale (right, horizontal comparison lines). *, **, *** indicate $p < 0.05$, 0.01, and 0.001 for the corresponding circular shift or session-swap tests, respectively.

### Figure S7

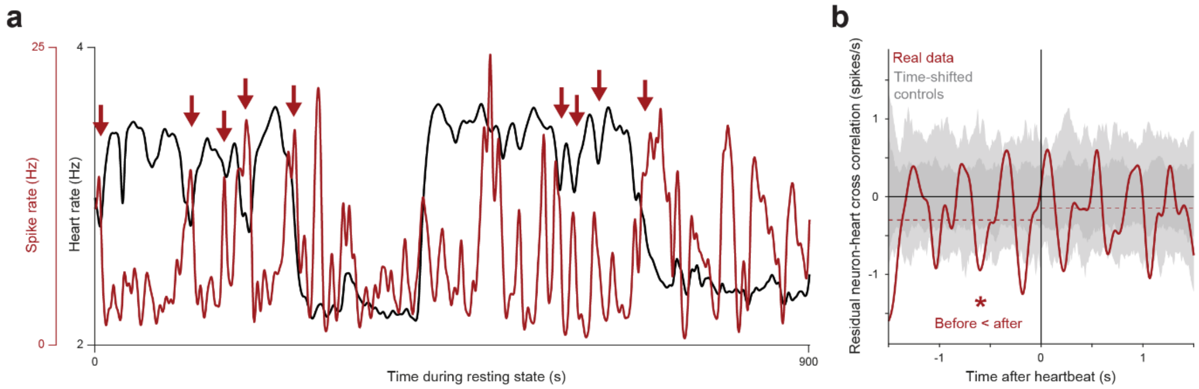


**Figure S7: Supplemental analysis illustrating the spike-heart order index. a,** An example neuron's firing rate (red) and heart rate (black) from the same session (same format as main text). This neuron appeared to have changes in activity negatively related to changes in heart rate, with sharp peaks in activity often occurring at the same time as, or slightly before, sharp reductions in heart rate (red arrows). **b,** Residual cross-correlation between spike rate and heart rate, expressed as residual firing rate aligned on the heartbeat (red; after regressing out linear, quadratic, and cubic effects of time; Methods). Shaded area shows the central 68% (dark gray) and 95% (light gray) of the same quantity computed from n=100 circularly time-shifted control datasets (Methods). The cross-correlation is overall shifted negatively when considering the full +/- 1.5 s extent of the cross-correlogram (i.e. many more data points are negative than positive), indicating an overall negative relationship between this neuron's spiking and heart rate over the timescale of seconds. In addition, the mean of the cross-correlation is more negative before than after the heartbeat (red dashed lines; left < right, $p < 0.05$, circular shift test), indicating that this neuron's negative changes in activity related to the heart rate tended to occur more before than after heartbeats. Thus, this neuron had a significant positive normalized spike-heart order index: its sign of activity modulation (negative) is aligned with the sign of its before-after difference in activity (before < after).

### Figure S8

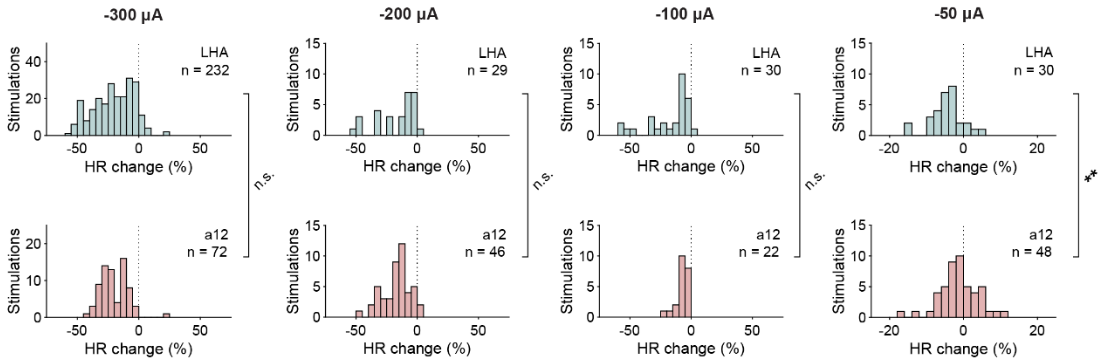


**Figure S8: Supplementary analyses comparing effective-site electrical stimulation-evoked responses in LHA and a12.** Histogram of the heart rate change effects averaged 0-1.5s after stimulation onset shown for -300, -200, -100, and -50 µA current. Top: Effective site stimulations in LHA. Bottom: Effective site stimulations in a12. ** next to the brackets indicates p < 0.01, n.s. means non-significant (rank-sum test).

### Figure S9

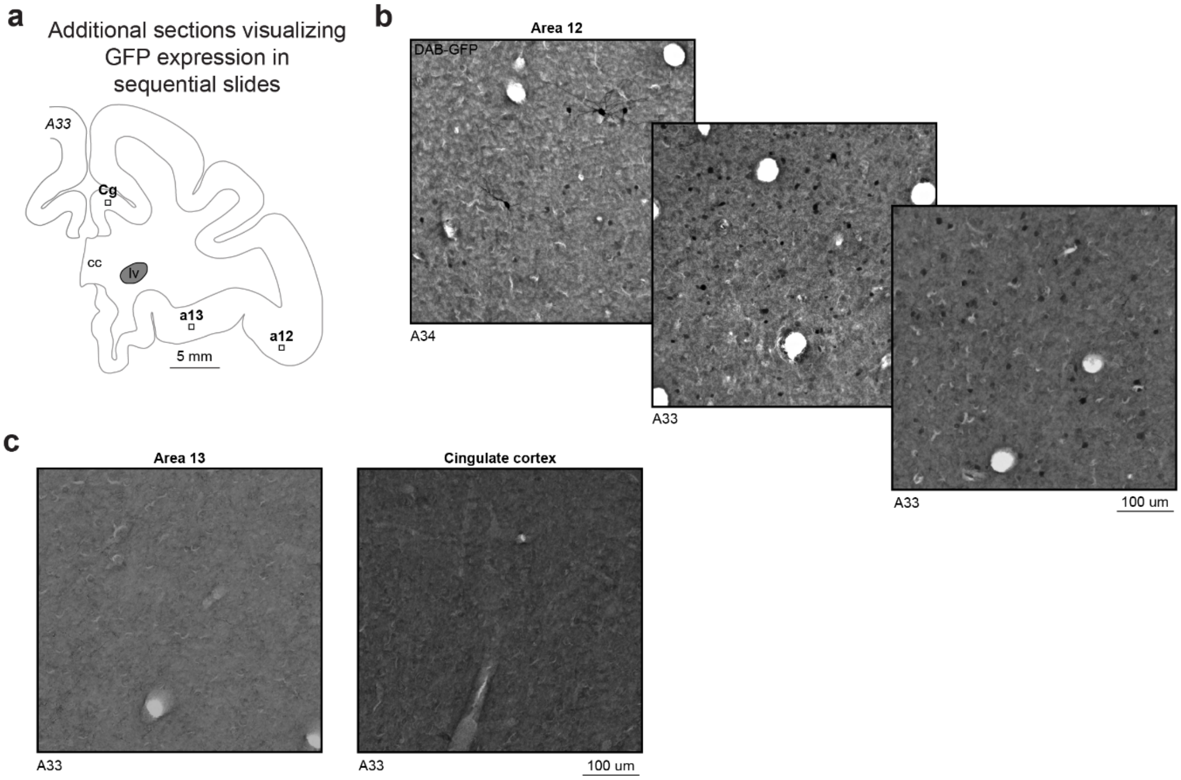


**Figure S9: Supplementary histology for chemogenetic experiments. a,** Schematic illustration of monkey brain in the coronal plane at interaural A33. Boxes indicate estimated imaging locations for a12, Cg, and a13. **b,** Representative IHC images visualizing DAB-stained GFP in area 12 in sequential slides. **c,** Representative IHC images visualizing DAB-staining in area 13 and cingulate cortex on the same slide with GFP-positive staining in area 12.

### Figure S10

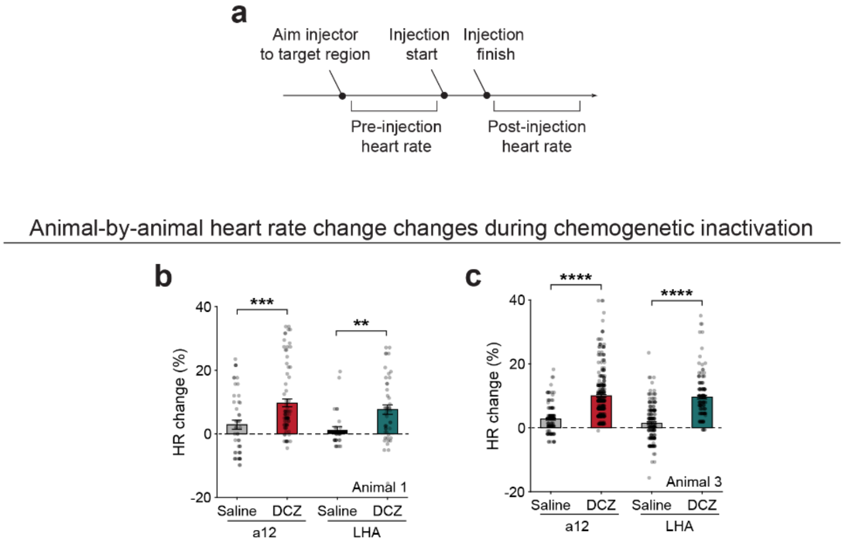


**Figure S10: Supplementary information for chemogenetic inactivation of a12→LHA. a,** Experimental timeline for acquiring data from chemogenetic inactivation experiment, which acquires both pre- and post-injection heart rate to normalize heart rate changes on a session-by-session basis. **b,** Heart rate changes during chemogenetic inactivation for animal 1. Top: Average heart rate change following saline or DCZ administration in a12 or LHA. Bottom: Histograms of the heart rate changes for each injection agent and injection location. Error bars represent s.e.m. **, *** above the brackets indicates $p < 0.01, 0.001$, respectively (rank-sum test). **c,** Heart rate changes during chemogenetic inactivation for animal 3. Top and bottom panels follow the same format as (b). **** above the brackets indicates $p < 0.0001$ (rank-sum test).

### Figure S11

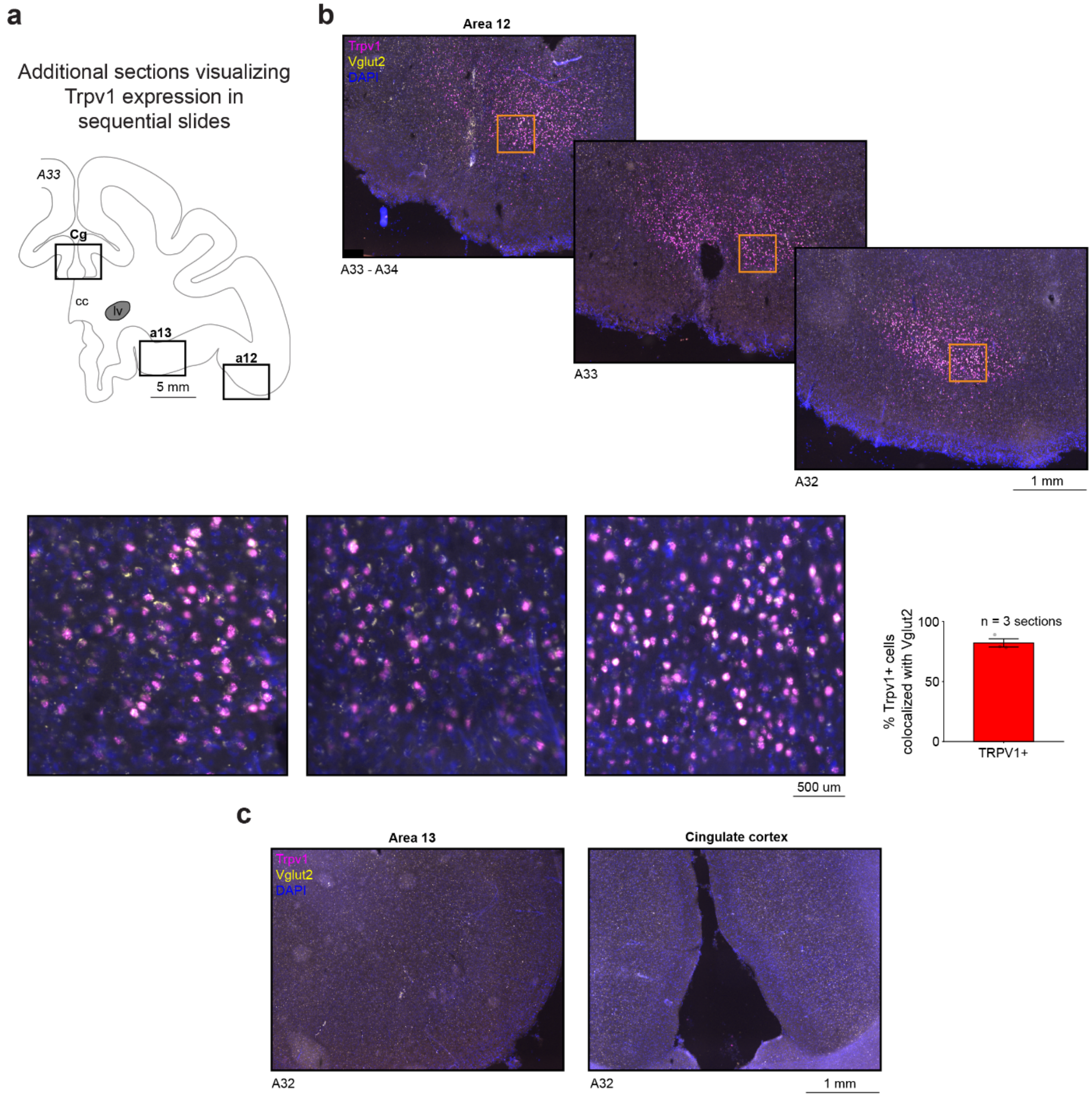


**Figure S11: Supplementary histology for sonogenetic experiments. a,** Schematic illustration of monkey brain in the coronal plane at interaural A33. Boxes indicate estimated imaging locations for a12, Cg, and a13. **b,** Top: Representative ISH images visualizing DAPI (blue), Trpv1 RNA (magenta), and Vglut2 RNA (yellow). Yellow boxes indicate regions of interest that are zoomed in on the bottom panels. Bottom, left: Representative zoomed in ISH images. Bottom, right: Quantification of cell-type specificity from injection of AAV encoding TRPV1 under the CaMKII promoter. **c,** Representative ISH images in area 13 and cingulate cortex on the same slide with Trpv1-positive staining in area 12.

### Figure S12

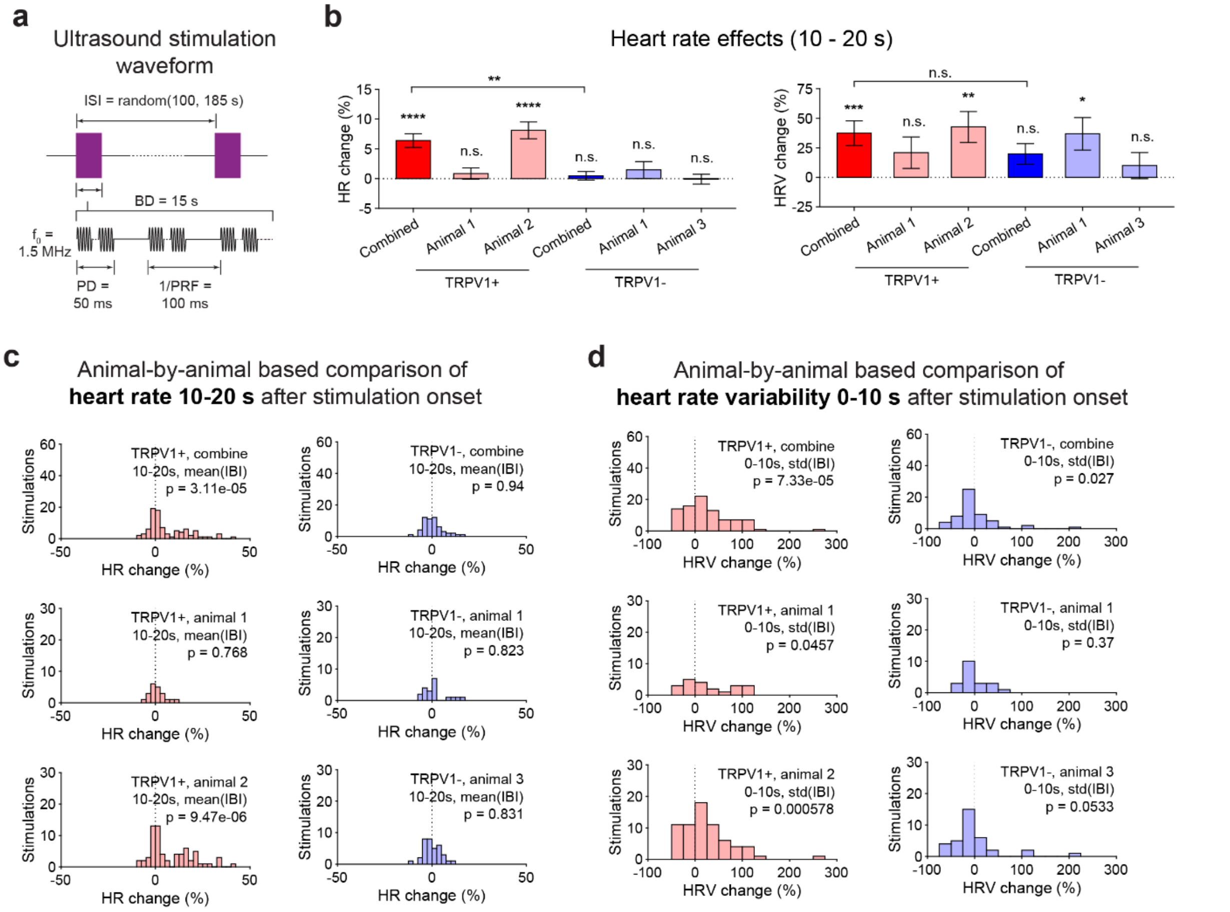


**Figure S12: Supplementary information for sonogenetic perturbation of a12. a,** Schematic illustration of the ultrasound stimulation waveform. **b,** Heart rate responses 10-20s after stimulation onset for TRPV1+ and TRPV1- animals, both combined and on an animal-by-animal basis. Left: Average heart rate responses. Right: Average heart rate variability responses. Error bars represent s.e.m. Significance above each bar corresponds to a one-sample signed-rank test against a hypothetical median of 0. Significance above the bracket indicates rank-sum test. **, **** indicates $p < 0.01$, 0.0001, respectively. n.s. means non-significant. **c,** Histograms of the average heart rate change 10-20s after stimulation onset. Left: histograms of the heart rate change for the TRPV1+ group with combined monkeys, animal 1, and animal 2. Right: Histograms of the heart rate change for the TRPV1- group with combined monkeys, animal 1, and animal 3. **d,** Histograms of the average heart rate variability change 10-20s after stimulation onset. Left and right panels follow the same format as (c).

### Figure S13

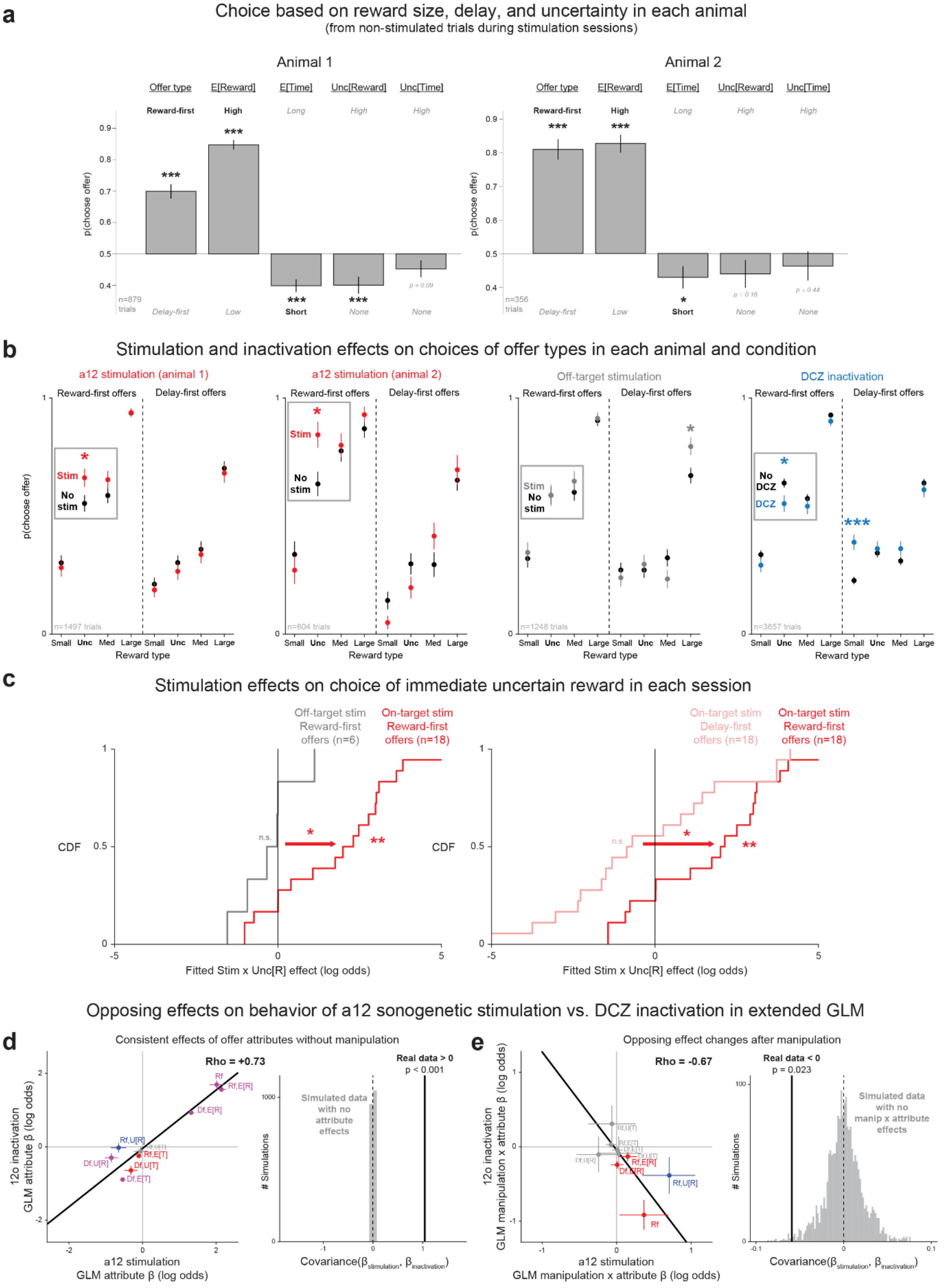

a
Choice based on reward size, delay, and uncertainty in each animal
(from non-stimulated trials during stimulation sessions)
Animal 1
Animal 2
Offer type
E[Reward]
E[Time]
Unc[Reward]
Unc[Time]
Reward-first
High
Long
Delay-first
Low
Short
None
p(choose offer)
n=879 trials
n=356 trials
p = 0.09
p = 0.16
p = 0.44
b
Stimulation and inactivation effects on choices of offer types in each animal and condition
a12 stimulation (animal 1)
a12 stimulation (animal 2)
Off-target stimulation
DCZ inactivation
Reward-first offers
Delay-first offers
Stim
No stim
No DCZ
DCZ
n=1497 trials
n=604 trials
n=1248 trials
n=3657 trials
Small
Unc
Med
Large
Reward type
c
Stimulation effects on choice of immediate uncertain reward in each session
Off-target stim Reward-first offers (n=6)
On-target stim Reward-first offers (n=18)
On-target stim Delay-first offers (n=18)
CDF
n.s.
Fitted Stim x Unc[R] effect (log odds)
Opposing effects on behavior of a12 sonogenetic stimulation vs. DCZ inactivation in extended GLM
d
Consistent effects of offer attributes without manipulation
Rho = +0.73
12o inactivation GLM attribute β (log odds)
a12 stimulation GLM attribute β (log odds)
Real data > 0
p < 0.001
Simulated data with no attribute effects
### Simulations
Covariance(β_stimulation, β_inactivation)
e
Opposing effect changes after manipulation
Rho = -0.67
12o inactivation GLM manipulation x attribute β (log odds)
a12 stimulation GLM manipulation x attribute β (log odds)
Real data < 0
p = 0.023
Simulated data with no manip x attribute effects

**Figure S13: Supplemental analysis demonstrating behavioral task performance and its modulation by sonogenetic and chemogenetic manipulations of a12. a,** Simple omnibus behavioral preference measures for each animal, based on non-stimulated trials during stimulation sessions. Same format as main text. **b,** Bidirectional manipulation effects on percent choice of offers based on offer type (reward-first or delay-first) and reward type (x-axis), separately for a12 stimulation in animals 1 and 2 (left, red), for off-target stimulation (middle, gray), and a12 DCZ inactivation (blue, right). Same format as main text. Consistent with our main results, in all cases, the choice of reward-first uncertain reward offers (gray boxes) was significantly increased by a12 stimulation, not significantly changed by off-target stimulation, and significantly reduced by DCZ inactivation; while no significant changes were observed in reward-first certain-reward offers or delay-first uncertain reward offers. DCZ also significantly altered choice of delay-first non-rewarded offers. **c,** Stimulation effects on choice were robust when examined on a session-by-session basis. We fitted the models to single session data using standard ridge regression as a form of regularization to obtain stable fits. Left: on-target vs. off-target stimulation. CDFs of per-session effects show a significant positive effect of on-target stimulation (red, $p < 0.01$, signed-rank test), no significant effect of off-target stimulation (gray) and a significant difference between them ($p < 0.05$, rank-sum test). Here, each session was fit using the model identified by the stepwise GLM (i.e. where stimulation interacts with motivation solely through a Stim x Reward-first Unc[R] effect). Right: on-target stimulation effects on reward-first vs. delay-first offers. CDFs of per-session effects show a significant positive effect of on-target stimulation on reward-first offers (red, $p < 0.05$, signed-rank test), no significant effect on delay-first offers (pink), and a significant difference between them ($p < 0.05$, signed-rank test). Here, each session was fit using the model identified by the stepwise GLM but augmented with an additional Stim x Delay-first Unc[R] effect (pink) for comparison as a control. **d-e,** Confirmation of opposing effects of a12 sonogenetic stimulation vs DCZ inactivation using an extended GLM with all possible interaction effects. We performed this analysis as an additional control to ensure that our results were robust to the particular choice of which manipulation x motivation term to use to quantify the behavioral effects (e.g. by contrast to the specific Manip x reward-first Unc[R] regressor identified by the stepwise GLM). Note that any significant finding in this analysis is especially noteworthy, because it has markedly lower power than our main text analysis, due to extending the model to include 8 extra regressors that had been rejected by the stepwise GLM because they did not significantly improve the fit. Therefore, any individual regressor is less likely to be detectable as significant, but we reasoned that we might still make inferences based on the overall patterns of fitted regression weights for stimulation vs. inactivation. **d,** Confirmation of consistent baseline behavior, indicated by consistent fitted main effects of offer attributes without manipulation. Left: positive relationship between fitted main effects of each attribute on stimulation sessions (x-axis) vs. inactivation session dataset (y-axis). Black line indicates a linear fit (type 2 regression) and text indicates the rank correlation. Error bars are +/- 1 SE. Colors indicate significance (blue: x-axis; red: y-axis; purple: both; $p < 0.05$, t-tests). Labels indicate attributes ("Rf": binary regressor set to 1 for reward-first offers and 0 for delay-first offers; "Rf, X": attribute X for reward-first offers; "Df, X": attribute X for delay-first offers). Right: quantification and statistical test of this relationship in terms of the covariance of the regression weights from stimulation vs. inactivation sessions. The covariance is strongly positive for the real data (black line), which is significantly more positive

($p < 0.001$) than the distribution of n=2000 stimulated datasets generated from the regression model by setting all main effects to 0, to represent the null hypothesis that there were no true attribute effects (gray). **e,** Confirmation of opposing manipulation effects, indicated by a negative relationship between the fitted interaction effects. Left: negative relationship between stimulation x attribute interaction effects (x-axis) vs. inactivation x attribute interaction effects (y-axis). Right: Significant negative covariance of the interaction effects ($p < 0.05$, black) compared to n=2000 simulated datasets generated from the regression model by setting all interaction effects to 0, to represent the null hypothesis that there were no true interaction effects (gray).

### Figure S14

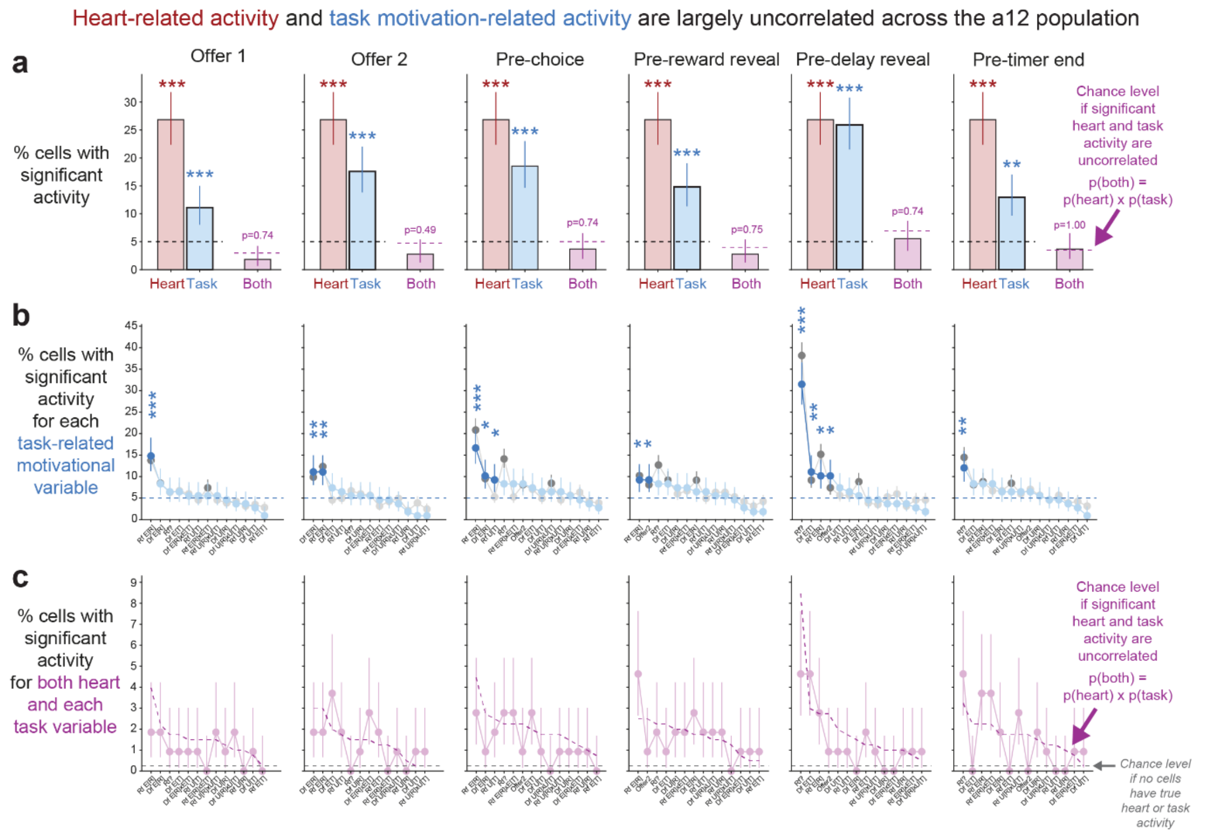


**Figure S14: Supplemental analysis demonstrating rest period heart-related activity and task period motivation-related activity are largely uncorrelated across the a12 population.** **a,** Significant heart- and motivation-related activity overlap at near- or below-chance levels in all task epochs. This is consistent with the hypothesis that these signals are “uncorrelated” or slightly “distinct” from each other, and is inconsistent with the hypothesis that these signals are “shared” across the population. Histograms show the percentage of neurons with significant heart-related activity measured using resting state data (red bar, same data in all columns; $p < 0.05$, circular shift test), significant motivation-related activity measured using task data (blue bar, $p < 0.05$, chi-squared test) in each of six task epochs (columns; Methods), and both heart and task significant (purple). Horizontal black dashed line indicates chance level for each signal alone (0.05). Horizontal purple line indicates chance level for overlap under the null hypothesis that significance of each significant signal occurs in the same number of neurons observed in the real dataset, but that heart and motivation are statistically independent of each other (i.e. p(both) = p(heart) x p(task)). Error bars are +/- 1 SE. In all task epochs, the percentage of cells with both signals significant is at or below chance level, with no significant difference from chance (all $p > 0.4$, exact binomial test of proportions). **b-c,** Similar results occur when considering each individual motivational attribute in the model. **b,** Percent of cells with significant effects of each individual attribute ($p < 0.05$, t-test) in each epoch (columns), sorted in descending order. Blue curve: all neurons with data recorded during both rest and task (n=108).

Gray curve: all neurons with data recorded during the task, regardless of whether they were recorded during rest (n=283), which produced similar results. Blue dashed line indicates the chance level (5%). Dark color indicates that more neurons are significant than expected by chance (*, **, *** indicate $p < 0.05$, 0.01, 0.001; exact binomial test). Labels on x-axis indicate the attribute names (“Rf?”: binary regressor set to 1 for reward-first offers and 0 for delay-first offers; “Rf, X”: attribute X for reward-first offers; “Df, X”: attribute X for delay-first offers; “Offer2”: binary regressor set to 1 if offer 2 was chosen and 0 otherwise). **c,** Percent of cells with significant effects of both heart-related activity and the corresponding motivational attribute (sorted in the same order as panel b). Gray dashed line indicates chance level under the simple null hypothesis that no cells have any true heart- or motivation-related activity (5% x 5% = 0.25%). Almost all data points exceed the gray line, indicating that significant heart- and task-related activity co-occur much more often than would be expected by chance if neurons had no true signals at all. Purple dashed line indicates chance level under the null hypothesis that significant heart- and task-related activity occur at the same levels observed in the real data but are statistically independent of each other (p(both) = p(heart) x p(task)). All attributes in all task epochs were similar to the null hypothesis of independence between heart- and task-related activity, with no significant differences (all $p > 0.05$).

**Table 1: Subject-by-experiment table.** Table showing the animal allocations across key experiments.

| Experiment | Animal 1 | Animal 2 | Animal 3 |
|---|---|---|---|
| Electrical stimulation in LHA | ✓ | ✓ | |
| Electrical stimulation in a12 | ✓ | | ✓ |
| Electrophysiological recordings with neural correlation | ✓ | ✓ | |
| Chemogenetic DREADD manipulation | ✓ | | ✓ |
| Sonogenetics TRPV1+ group | ✓ | ✓ | |
| Sonogenetics TRPV1- group | ✓ | | ✓ |
| Task behavior | ✓ | ✓ | |
